# Imaging the ultrastructures and dynamics of live erythrocyte membranes at the single-molecule level with a far-red probe on a microfluidic platform

**DOI:** 10.1101/2021.04.22.441044

**Authors:** Zhiwei Ye, Wei Yang, Shujing Wang, Ying Zheng, Xiaodong Zhang, Haibo Yu, Chunxiong Luo, Xiaojun Peng, Yi Xiao

## Abstract

Both the ultrastructures and dynamics of living erythrocyte membranes provide critical criteria for clinical diagnostics. However, it is challenging to simultaneously visualize these features at the single-molecule level due to the rigid photophysical requirements of different single-molecule imaging techniques. Herein, we rationally developed a far-red boron dipyrromethene membrane (BDP-Mem) probe that not only retained consistent and intensive single-molecule emission but also possessed the capability to photoswitch on cellular membranes. We also constructed a microfluidic platform for the noninvasive trapping and long-term imaging of nonadherent erythrocytes. By combining these advantageous technologies, super-resolution reconstruction and single-molecule tracking of living human RBC membranes were achieved at the molecular scale in a high-throughput fashion. Our integrated paradigm defines a quantitative approach for analyzing living RBC membranes under physiological and pathological conditions, improving imaging precisions and revealing new perspectives for future disease diagnostic approaches.

## Introduction

The analyses of erythrocytes (red blood cells, RBCs) provide diagnostics used to determine an individual’s health status. Two of the most significant RBC indicators are the ultrastructures and dynamics of their membrane envelopes.^1–3^ Nanoscopic imaging^4,5^ of membrane details is a powerful approach for improving diagnostic precision and dimensionality. However, these fine membrane architectures remain hidden due to the challenges involved in deploying living erythrocytes for singlemolecule imaging techniques. The first limiting issue includes the severe autofluorescence of hemoglobin-rich RBCs (Supplementary Fig.1) as well as the photophysical requirements of single-molecule techniques,^6–8^ which call for the development of high-performance probes in the long-wavelength region. The other limiting issue is the drifting of nonadherent RBCs during image acquisitions (Supplementary Fig.2, Supplementary Video 1), which requires the construction of exquisite platforms for noninvasive long-term imaging.

In this study, we integrated multidisciplinary techniques for simultaneously superresolving the ultrastructures and dynamics of living RBC membranes by combining a far-red boron dipyrromethene probe (BDP-Mem) with a versatile microfluidic device. Membrane-specific BDP-Mem was rationally constructed to have high brightness, a prolonged bright state and the capability to photoswitch in a hydrophobic microenvironment, fitting the demands of both single-molecule localization and single-molecule tracking. A microfluidic platform was developed to immobilize living RBCs in a long-term fashion, allowing super-resolution imaging under total internal reflection fluorescence (TIRF) field without shifting. With the precise manipulation of microenvironments, this platform was further capable of building artificial disease models. Through the fusion of these two advancements, both the ultrastructures and dynamics of living RBC membranes were superresolved with robust integrity at the molecular scale in a high-throughput fashion, providing multidimensional indicators for the RBC diagnostics.

## Results

### Far-red probe

Single-molecule localization and tracking constitute approaches for uncovering the ultrastructures and dynamics of RBC membranes. However, these two techniques have different fluorescent probe requirements. Although high brightness is needed for both techniques, the precondition for single-molecule localization is photoswitching,^8^ while that for single-molecule tracking is consistent single-molecule emission.^9–11^ It is a great challenge to construct fluorophores that satisfy both of these terms. Specifically, the prolongation of the bright state of a fluorophore considerably enhances the likelihood of bleaching, which forbids its recovery to the bright state and makes it more difficult to achieve photoswitching, especially without an oxygen-scavenged imaging enhancing buffer.

To determine a solution for this dilemma, we rationally developed a membrane probe based on the modern boron dipyrromethene (BODIPY) fluorophore. The parent BODIPY fluorophore exhibits high brightness and photostability in the hydrophobic environment found in cell membranes,^12,13^ and is eligible for single-molecule tracking. However, without an imaging enhancing buffer, BODIPY does not exhibit sufficient photoswitching performance for localization-based super-resolution imaging.^14^ Here, the core of BODIPY is extended with *p*-piperazinyl styryl units, the decoration of which enables the possibility of darkening through cis-trans isomerism after laser depletion.^15^ Such an enlarged conjugation also shifts the excitation and emission wavelength into the far-red region, successfully avoiding the autofluorescence from RBC hemoglobin. The quaternization of the piperazinyl groups further enhances the brightness by inhibiting nonradiative decay, according to our previously reported twisted intramolecular charge transfer (TICT) inhibition strategy.^16^ Moreover, the two positively charged moieties act as hydrophilic terminals, and together with the hydrophobic conjugation skeleton, endow the probe with amphipathic characteristics for targeting cell membranes. Therefore, an efficacious far-red probe, BDP-Mem (Fig.1a), which has photoswitching and high photostability characteristics, is designed to stain the membranes of RBCs for single-molecule correlated imaging studies.

**Fig.1.**
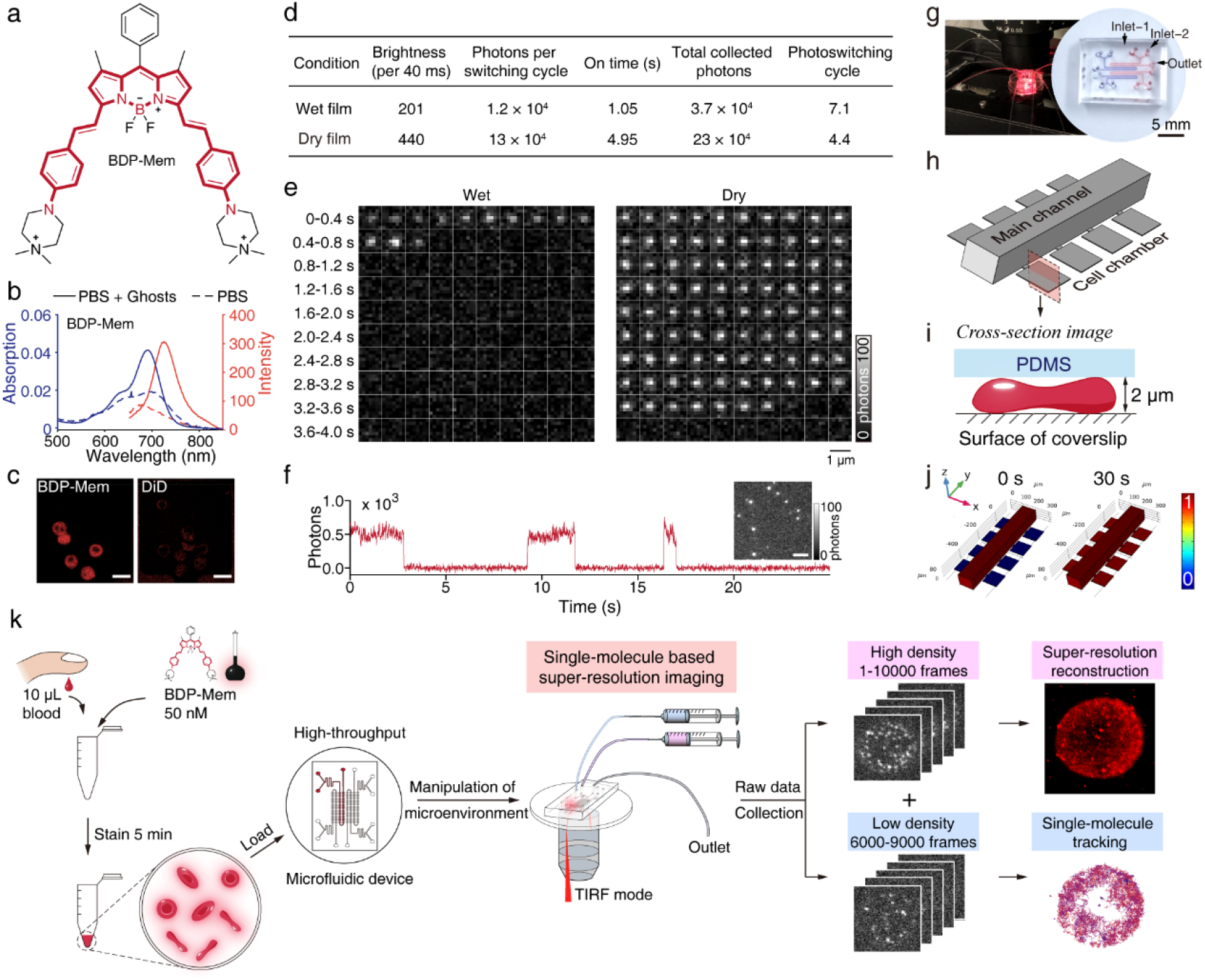
Construction of a far-red probe and a microfluidic device for imaging erythrocyte membranes. (a) Molecular structure of BDP-Mem. (b) Absorption and fluorescence spectra of BDP-Mem upon staining RBC Ghosts. (c) Confocal fluorescent images of RBCs stained with BDP-Mem or DiD under identical conditions. (d) Summarized single-molecule characteristics of BDP-Mem. (e) Time-series stack images of typical BDP-Mem molecules on a wet or dry film. (f) A single-molecule intensity trajectory of BDP-Mem on a dry film. Inset shows dispersed single-molecule signals. (g) Implementation of a microfluidic chip for imaging. An example of microfluidic chip filled with red or blue ink is shown on the right. (h) Schematic chart depicts the geometry of one channel. (i) Cross-section view of a cell chamber. (j) Comsol simulation of the time evolution of buffer exchange on the microfluidic device. (k) Schematic illustration of our approach for imaging RBCs membranes. Scale bars: 10 μm (c); 3 μm (f).

Spectral measurements reveal the robust brightness of BDP-Mem in the far-red region (Supplementary Fig.3 and Supplementary Table 1). BDP-Mem consistently exhibits far-red fluorescence in both protonic and nonprotonic solvents, displaying fluorescence peaks at 671-727 nm and the corresponding absorption peaks at 650-677 nm. This probe further demonstrates high molar absorption coefficients (ε= 9.8~10 × 10^4^ mol·L^−1^·cm^−1^) and quantum yields (0.52~0.54) in moderately polarized solvents (dichloromethane and methanol), which facilitates the staining of membranes with hydrophobic interiors. The excellent membrane-staining functionality of BDP-Mem was confirmed through the 3-fold fluorescence enhancement that occurred upon staining RBC ghosts (Fig.1b). Such enhancement is in sharp contrast with the minor changes that were induced by the commercially available membrane probes DiI and DiD (Supplementary Fig.4). Furthermore, on living RBC membranes, BDP-Mem demonstrated an 8-fold increase in fluorescence (Fig.1c, Supplementary Fig.5) and enhanced photostability (Supplementary Fig.6) compared with DiD. These results validate the advantages of this probe over commercial probes for imaging RBC membranes.

### Single-molecule properties of BDP-Mem

The single-molecule performance of BDP-Mem was further investigated on a polyvinyl acetate (PVA) film. Dry films are useful for mimicking hydrophobic membranes, and wet films, formed in humid environment, exhibit characteristics similar to hydrophilic media. BDP-Mem demonstrates enhanced brightness (2.2-fold) and prolonged on time (the duration of bright state, 4.7-fold) on a dry film compared to that on a wet film (Fig.1d). This sharp contrast is further displayed in time-series stack images (Fig.1e) of the single-molecule signals from BDP-Mem. Consequently, under hydrophobic conditions, BDP-Mem exhibits a more than 10-fold enhancement of the photon collection per switching cycle (13 × 10^4^ dry *vs*. 1.2 × 10^4^ wet; roughly equal to the product of on time and single-molecule brightness) and a 6-fold enlargement in the number of total collected photons (23 × 10^4^ dry *vs*. 3.7 × 10^4^ wet) before photobleaching. The bright and stable BDP-Mem signals are preferable for long-term single-molecule tracking on membranes. Moreover, BDP-Mem demonstrates profound photoswitching (Fig.1f; average cycle numbers: 4.4 dry and 7.1 wet) on the films, which is beneficial for obtaining high localization density in super-resolution reconstruction. Additionally, on living membranes, the motion of our probe during a prolonged bright state continually provides additional localization data for plotting the details of the geometric surface. In general, our probe exhibits preferable single-molecule photophysical properties, displaying photoswitching for super-resolution imaging and prolonged on time for single-molecule tracking.

### Microfluidic imaging platform

The high signal-to-noise ratio required for high-quality super-resolution imaging is achieved through a thin illumination layer of the TIRF field, which requires the immobilization of nonadherent RBCs to a TIRF depth of 200 nm. However, chemical fixation through methanol or polylysine irreversibly changes the cellular conditions, leading to incredible results for living-cell studies.^17^ Microfluidics has emerged as an alternative and effective approach for long-term studies of suspended cells. Through this technique, *E. coli*,^18^ yeast^19^ and red blood cells^20^ have been spatially restricted and investigated in micron-sized chambers and other geometries. Moreover, microfluidics has been shown to enable precise manipulation of the cell microenvironment at a high spatiotemporal resolution,^21^ making it an ideal tool to integrate into the study of RBC super-resolution imaging.

Our microfluidic platform is shown in Fig.1g. This device has four parallel main channels with an array of 45 chambers (100 μm × 100 μm × 2 μm) connected to each side of these channels (Fig.1h). The height of each chamber was designed to be the same as the height of a human red blood cell, to horizontally confine the loaded cells for imaging within the TIRF depth (Fig.1i). Benefitting from the micro size of the device, the sample loading of the chip was completed with only 10 μL of whole blood. Through pumping stimulus into the main channel at the additional inlet, the microfluidic platform further allows studies to be conducted under physiological and pathological conditions. Under simulation (Multiphysics, Fig.1j), microenvironment exchange within the cell-chambers achieves equilibrium in 30 s, which is rapid compared to the cellular response, preventing biased results from insufficient replacement.

Our combined approach for imaging erythrocytes is demonstrated in Fig.1k. Fingertip blood (10 μL) was stained with BDP-Mem (50 nM) for 5 min in anticoagulant media. After direct loading into the microfluidic device, RBCs were autonomously self-isolated into microchambers. To construct artificial disease models, the microenvironments for RBCs in cell chambers were actively controlled to mimic physiological and pathological conditions. Then, 10000 stack frames with single-molecule signals were collected with a TIRF microscope. All frames were utilized for localization to obtain high-density super-resolution reconstruction. The frames with low-density single-molecule signals (6000-9000 frames) were applied to singlemolecule tracking studies. Therefore, through the combination of our probe and microfluidic device, it is practical to simultaneously investigate the ultrastructures and dynamics of living RBC membranes.

### Super-resolution imaging of a living erythrocyte membrane

Live erythrocyte membranes were super-resolution imaged through our approach. BDP-Mem exhibits excellent photoswitching performance (Supplementary Video 2) on the membrane of a live RBC. Remarkably, this blinking of BDP-Mem on membranes generates in normal culture media regardless of cell types (as evidenced by the result on a HeLa cell in Supplementary Fig.7), which prevents cellular damage caused by the addition of oxygen-depletion reagents and any other imaging enhancing buffer. In addition, the small volume of each microchamber (20 pL) allows fluorescent interference from free fluorophores in the medium to be distinctly avoided, resulting in a clean background. Therefore, our method enables high-quality super-resolution imaging of a live RBC membrane (Fig.2a), in stark contrast to the conventional imaging result (Figs.2b-c). The specialized biconcave RBC morphology under the TIRF mode is precisely depicted through super-resolution reconstruction with an average localization uncertainty of 23.2 nm (Fig.2d). The spatial resolution is further estimated by a Fourier ring correlation (FRC) analysis (Fig.2e),^22^ revealing a reconstruction resolution of 68.5 nm. The high-quality super-resolution reconstruction of BDP-Mem is attributed to its significantly enhanced single-molecule brightness (Fig.2f) and excessive resistance to photobleaching (Fig.2g). Contrary to the superior results from our probe, reconstruction conducted via the commercial membrane probe DiD fails to plot the morphology of a RBC membrane because there are only a few localization spots defined (Supplementary Fig.8).

**Fig.2.**
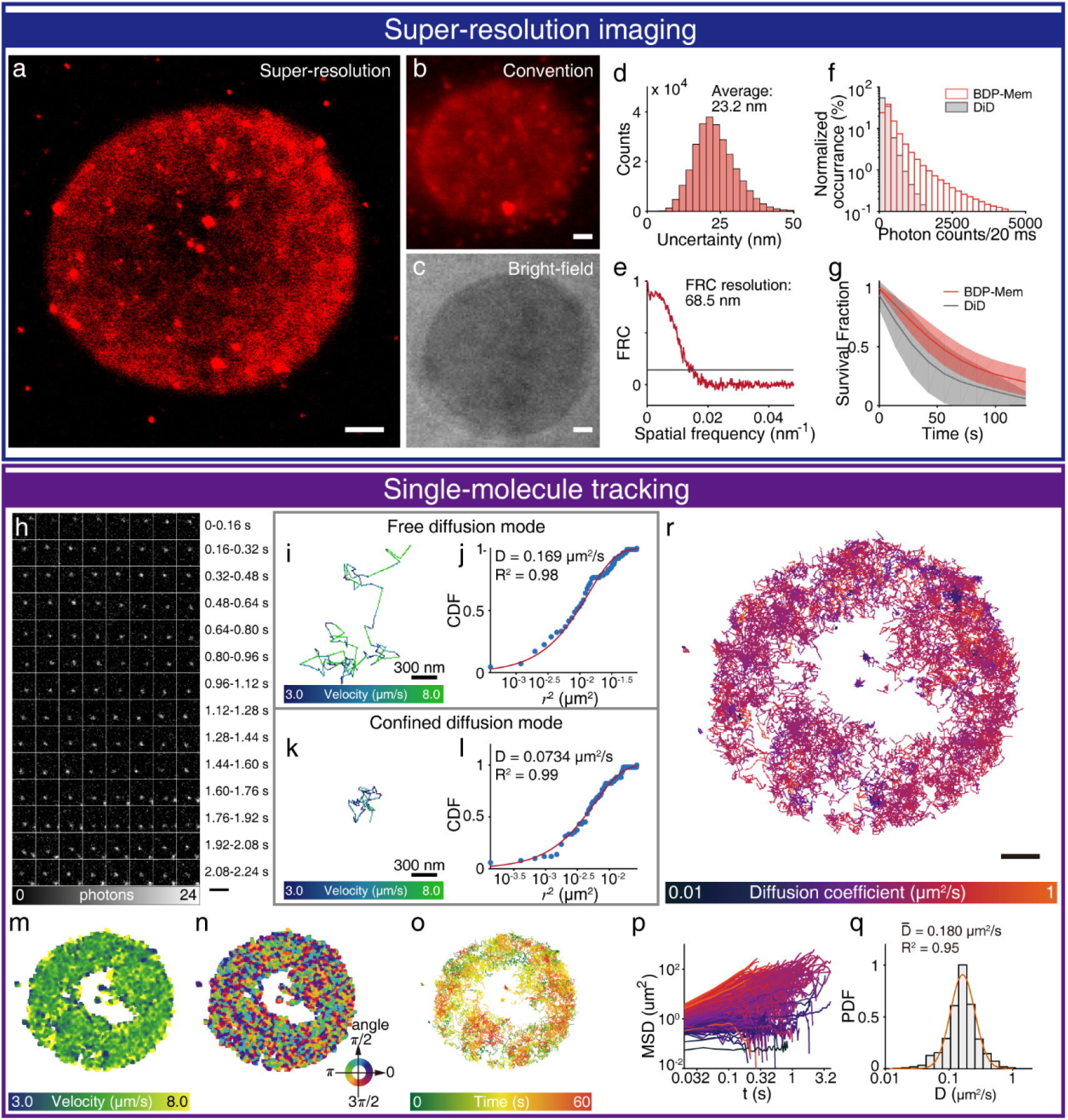
Imaging the ultrastructures and dynamics of a living RBC membrane at the single-molecule level through our combinational approach. (a) Localization of BDP-Mem reconstructs the membrane morphology of a red blood cell. The convention fluorescent image (b) and bright-field image (c) of the same cell. (d) Histogram of localization uncertainties. (e) Fourier ring correlation (FRC) analysis of the reconstruction. Comparison of single-molecule brightness (f) or survival fractions (g) between BDP-Mem and DiD (n > 10 RBCs) during super-resolution imaging. (h) Time-series stack images of a typical BDP-Mem molecule on the RBC membrane. (i,k) Magnified view of two typical molecular trajectories, with color indexed by their velocities. (j,l) The cumulative distribution function (CDF) of the square displacements from trajectories (i) and (k). The CDF was fitted with a single exponential function. The velocity map (m), direction map (n) and time-colored trajectory overlay (o) from the same RBC. (p) The mean square displacements (MSD) log plots of BDP-Mem. (q) The probability density function (PDF) of the diffusion coefficients, fitted with a log-normal distribution. (r) Overlay of single-molecule tracking trajectories (over 10 steps) of BDP-Mem on the membrane. Colors are indexed by their diffusion coefficients. Scale bars: 1 μm (a-c, r); 2 μm (h).

### Single-molecule tracking and diffusion analyses

The capabilities for long-term trapping of living RBCs inside our chip and the enduring fluorescence of BDP-Mem enable single-molecule tracking studies. The single-molecule signals on the RBC membrane existed for a period of time, which was beneficial for tracking and consistent with the dry film results. Fig.2h shows 2.24 s time-series stack images of a typical BDP-Mem molecule on the RBC membrane. The trajectory of this molecule is depicted and analyzed in Figs.2i-j, following the free diffusion mode. Another BDP-Mem molecule following the confined diffusion mode is depicted and analyzed in Figs.2k-l (more molecules are presented in Supplementary Fig.9). These molecules show heterogeneous diffusive speeds on the membrane, which indicates diffusive limitations for some loci of the membrane. This limitation might correlate to negative potentials. Further, this velocity discrepancy is accurately mapped on the membrane (Fig.2m) alongside the corresponding molecular movement directions (Fig.2n). The temporal distribution of tracked molecules were exhibited in a track overlay in Fig.2o, indexed with their acquisition time.

Moreover, the two modes of diffusion are revealed through the plots of the mean square displacements (MSDs, Fig.2p) of the molecules. The dark blue colored lines (with a flattened trends) represent molecules with confined diffusion, while the red colored lines (with an inclined trends) correspond to molecules under the free diffusion mode. To quantitatively measure molecular motion, the diffusion coefficient of a single molecule was calculated by fitting the cumulative distribution function (CDF) of its square displacements to a single exponential function, as shown in Figs.2j and l (R^2^ ≥ 0.98). Molecules following free diffusion modes exhibit apparent increased diffusion coefficients compared to those molecules following confined diffusion modes (e.g., 0.169 *vs* 0.0734 μm^2^/s). Then, the distributions of the diffusion coefficients are further analyzed. For most cells, since few molecules exist in the confined diffusion mode, the distributions are fitted well by a log-normal distribution (e.g., R^2^ = 0.95, Fig.2q). For RBCs with more confined molecules, the distributions present two log-normal distributions (e.g. Cell5 in Supplementary Fig.10). Finally, tracks with > 10 steps were overlaid in Fig.2r. The colors of the trajectories are indexed on the basis of their diffusion coefficients. The variance in color lightness reflects heterogeneity in the geometrical dynamics across the RBC membrane.

### RBC membranes under osmotic stress

We applied our method to visualize and understand the delicate transformations of RBC membranes under stomatocyte-discocyte-echinocyte sequences (Figs.3a-e) through the application of hypotonicity-isotonicity-hypertonicity osmotic pressures.^23^ Variation in blood osmolality commonly occurs with abnormal intakes of salt, nutrients or other drugs. To obtain different osmotic stress conditions, we adjusted the media osmolarity by the addition of salt or MilliQ water. In an isotonic buffer (310 mOsm/kg, Fig.3c), the RBCs preserved as biconcave discocytes. As the osmolarity decreased (251 mOsm/kg, Fig.3b), these cells gradually swelled to spheres, forming stomatocytes. When the osmolarity further decreased (143 mOsm/kg), these cells burst into leuco vesicles (Fig.3a), permanently losing their inner contents. The swelling deformation of RBCs is further evidenced by the flattened slope of the intensity profiles, from the edge to the center, during hypotonic development (Lines 1-3, Fig.3f). On the other hand, with increasing osmolarity (Figs.3c-e), RBCs transformed to a series of crenated echinocytes. In a less hypertonic solution (381 mOsm/kg, Fig.3d), the cell gradually shrank about its center, with a few wrinkles left around the middle of the cell. As the osmolarity further increased to 466 mOsm/kg, RBCs dramatically shrank, leaving protrusions at the TIRF depth (Fig.3e). The diameters of these protrusions were measured at approximately 100 nm (Lines 4-7, Fig.3f), in appreciable contrast to the rough estimations (>290 nm) that were obtained from the conventional image (Supplementary Fig.11). Therefore, we systematically investigated the morphological transformations of RBC membranes through single-molecule localization super-resolution imaging with our imaging platform.

**Fig.3.**
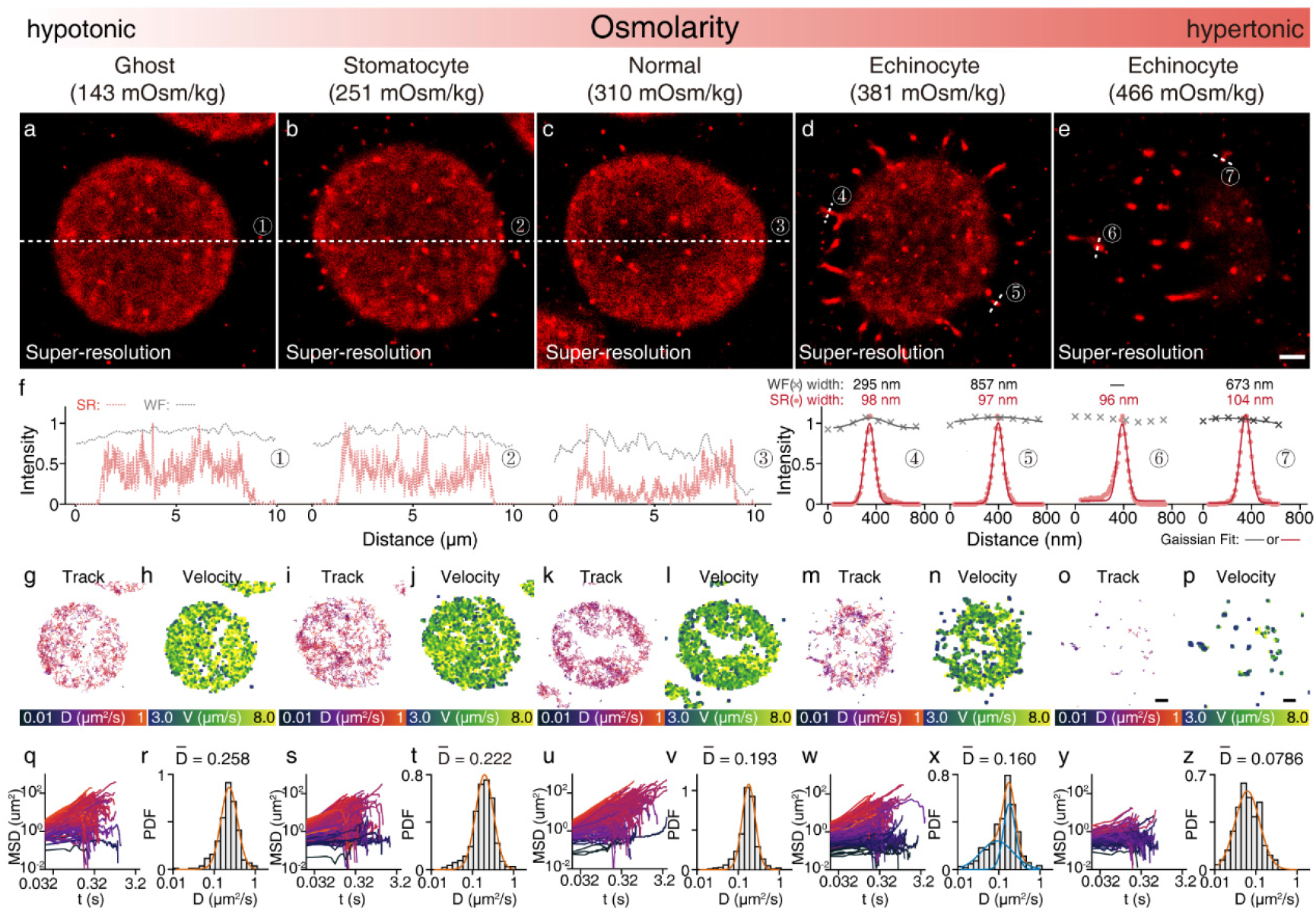
Ultrastructures and dynamics of RBC membranes under osmotic stresses. (a-e) Superresolution images of the membrane morphology from typical RBCs. (f) Comparison of the intensity profiles on the dashed lines highlightened in both super-resolution (a-e) and conventional images (Supplementary Fig.11). Single-molecule trajectory overlays (g, i, k, m, o), velocity maps (h, j, l, n, p), MSD log plots (q, s, u, w, y) and diffusion coefficient (D) distributions (r, t, v, x, z) of BDP-Mem on RBC membranes. Colors of MSD plots are indexed by diffusion coefficients, consistent with trajectory overlay images. Scale bars: 1 μm.

We further performed single-molecule tracking studies (Figs.3g-z) on the RBC membranes under osmotic pressures. The overlaid trajectories (Figs.3g, i, k, m, o) and the velocity maps (Figs.3h, j, l, n, p) show heterogeneous probe diffusion on the membranes. During the deformation from discocytes to echinocytes, the diffusion coefficient of BDP-Mem on the membranes sharply decreases (from 0.193 μm^2^/s to 0.160 μm^2^/s or 0.0786 μm^2^/s; Figs.3v, x, z). A consistent trend is shown in the MSD plots (Figs.3u, w, y), with the decrease of inclined components (free-diffusive molecules) following the increase of osmolarity. Under hypertonic conditions (381 mOsm/kg), the fraction of confined diffusive molecules enlarges, forming double lognormal distributions (Fig.3x). When hypertonicity is further lifted to 466 mOsm/kg, the probes are profoundly constrained on the RBC membrane, with few free-diffusive molecules. When the osmolarity decreases with the formation of stomatocytes and ghosts, the membrane dynamics increase, consistent with their swelled morphologies (0.222 μm^2^/s stomatocyte, 0.258 μm^2^/s ghost; Figs.3r, t). The above quantitative analysis reveals a consistency between the dynamics and morphological transformations of RBCs in that the diffusion coefficients monotonously decrease along with RBC shrinkage under increasing osmotic pressures.

### RBC membranes under lead stress conditions

In addition, we investigated the ultrastructures and dynamics of live erythrocyte membranes under lead stress conditions. Lead ion (Pb^2+^) is one of the most poisons to RBC membranes,^24^ since this ion massively accumulates in human RBCs at a ratio of approximately 95%.^25^ Our super-resolution imaging reveals the transformation of RBCs from discocytes to spiked echinocytes under Pb^2+^ stress conditions. Within 1 h of Pb^2+^ incubation, the erythrocytes gradually shrank with the appearance of protrusions on their edges due to dehydration (Figs.4a-f). When the incubation time was prolonged to 2 h, RBCs exhibited further crenation, turning into spiked echinocytes (Figs.4g-l). This transformation was inhibited by the addition of Senicapoc, which is a potential drug for the treatment of sickle cell disease and prevents dehydration by inhibiting Gardos channels. Under this condition, RBCs maintained their spherical shape even after 2 h of incubation with Pb^2+^ (Figs.4m-r, s-x). The dynamics of membrane are also investigated. After more than 1 h poisoning with Pb^2+^, although the morphology of RBCs severely transformed, the diffusion coefficient decreases marginally (0.222, 0.181 μm^2^/s, Pb^2+^; Figs.4f, l). When Senicapoc is added to block dehydration, the diffusion coefficient of BDP-Mem remains almost constant (0.228, 0.213 μm^2^/s Pb^2+^ + Senicapoc; Figs.4r, x). For acute Pb^2+^ conditions, the membrane dynamics demonstrate a different tendency toward morphological deformation, which is distinct from the membrane fluidity shifts under osmotic pressures. Therefore, in addition to the ultrastructures, the membrane dynamics are another important indicator for evaluating the physiological states of erythrocytes.

**Fig.4.**
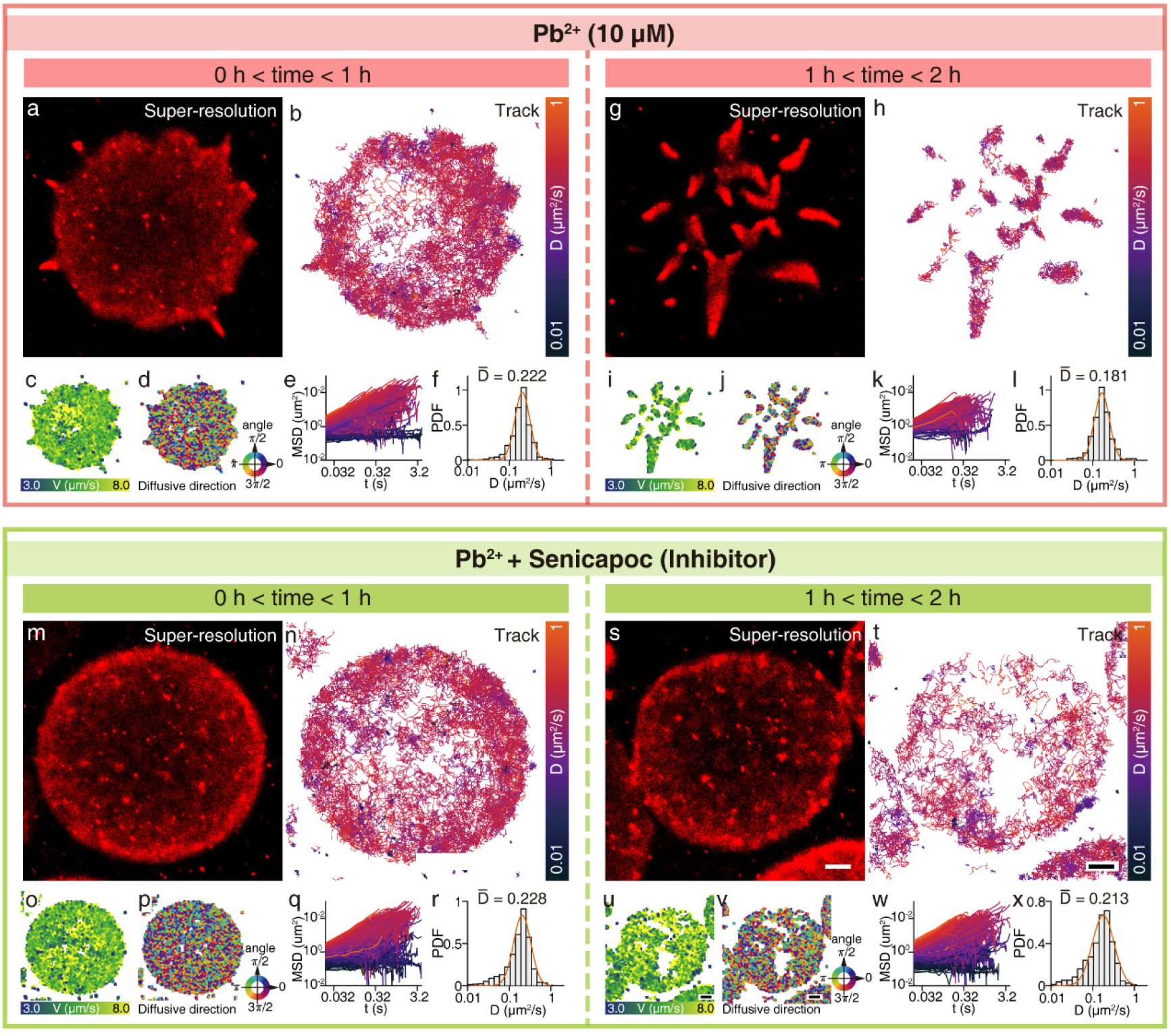
Ultrastructures and dynamics of living RBC membranes under Pb^2+^ poisoning stress conditions. RBCs were treated with Pb^2+^ stimulus (a-l) or in combination with Senicapoc (m-x). For each cell, the top row presents super-resolution reconstruction (a, g, m, s) and the single-molecule trajectory overlays (b, h, n, t); the bottom row presents the velocity maps (c, i, o, u), the direction maps (d, j, p, v), MSD log plots (e, k, q, w) and the diffusion coefficient distributions (f, l, r, x) of BDP-Mem on the RBC membranes. Scale bars: 1 μm.

### High-throughput analysis

Since our on-chip platform integrates 4 main channels, each containing 90 chambers, it further allows super-resolution imaging and single-molecule tracking studies of RBCs in a high-throughput fashion. Overall, we tested more than 140 RBCs. Among them, 50 RBCs under 10 different conditions were systematically analyzed and are displayed in Supplementary Fig.10. The collected results present consistent morphological and fluidic transformations related to the instances of RBC membranes discussed in the previous sections. Furthermore, the high-throughput results enable a statistical analysis on the fluidity of RBC membranes under various conditions (Fig.5). Under osmotic stress, the dynamics of RBC membranes significantly decrease with increasing osmolarity (Fig.5a, Supplementary Table 2), corresponding to their enormous morphological deformations. In contrast, under Pb^2+^ poisoning conditions, the diffusion coefficients exhibit no significant differences (Fig.5b, Supplementary Table 3), even though the RBC morphologies are unambiguously transformed. Membrane ultrastructures and dynamics are two significant indicators for both evaluating and diagnosing the states of RBCs. Under different conditions (osmotic or lead stress), RBCs demonstrate differentiated response characteristics. The combination of two indicators accurately depicts the states of RBCs in full, while a single indicator provides biased diagnostics. Our approach is feasible, high-throughput and multidimensional, simultaneously providing plentiful details about both the ultrastructures and dynamics of membranes, facilitating the precision improvement of RBC diagnostics.

**Fig.5.**
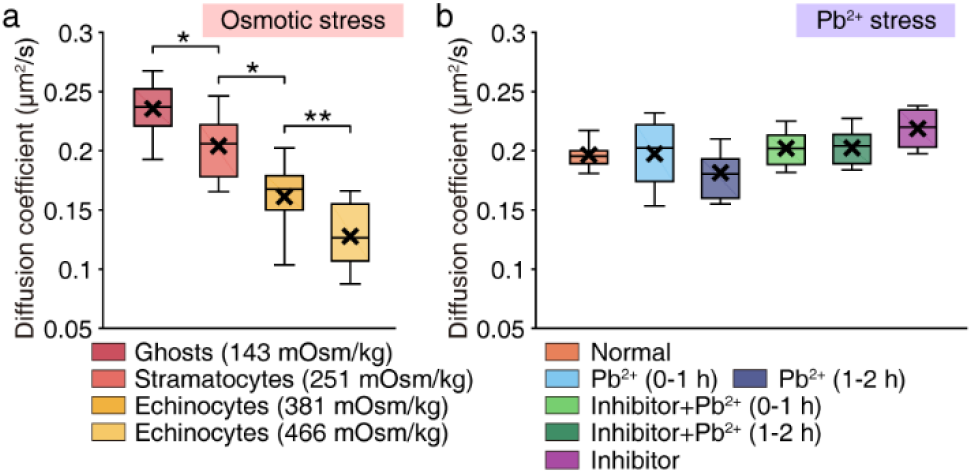
Comparison on the average diffusion coefficients of BDP-Mem on the RBC membranes under osmotic stress (a) and Pb^2+^ poisoning (b) conditions. The boxplot elements are defined as following: center line, median; box limits, upper and lower quartiles; whiskers, the 10th percentile and the 90th percentile. Significance was determined through a two-sample t-test with p < 0.05, *; < 0.001, ** (total number: n = 147 RBCs).

## Discussion

In summary, since ultrastructures and dynamics at the single-molecule level are two critical diagnostic indictors, we have integrated multidisciplinary approaches for live-cell super-resolution imaging of human RBC membranes to resolve these quantitative details. On one hand, a new membrane-specific probe, BDP-Mem, was rationally constructed to realize far-red emission that avoids the intense autofluorescence of RBCs. This probe further demonstrated high brightness, a prolonged bright state and photoswitching performance on cell membranes, qualifying as a versatile probe for both single-molecule localization and single-molecule tracking microscopies. On the other hand, a microfluidic device was designed for long-term noninvasive immobilization of living nonadherent RBCs. The on-chip study further enabled quantitative analyses of morphologically transformed RBCs under disease conditions in a high-throughput fashion. The collaboration of these two advancements defined multidimensional parameters for analyzing both morphological and dynamic transformation of RBC membranes at the nanoscale, providing new insights for both expanding the dimensions and enhancing the precision of disease diagnostics involving RBCs.

## Methods

### Super-resolution imaging

Super-resolution imaging experiments were performed on an Olympus IX71 inverted microscope with a ×100 objective (UAPON 100×OTIRF; 1.49 numerical aperture) as described earlier.^16,26^ Light from continuous 640 nm laser (100 W, Coherent) was controlled by an acousto-optic tunable filter (AOTF) and focused on the back focal plane of the objective. Evanescent lighting was achieved through a motorized TIRFM illuminator (IX2-RFAEVA-2, Olympus). Stack images was collected on a high-sensitive EMCCD (electron-multiplying charge-coupled device, iXon DU-897U) camera at an exposure time of 20 ms. Axial drift of stages (MS-2000, assembled with a top piezo-Z stage, PZ-2150FT, ASI) was corrected with a CRISP Autofocus system (ASI).

### Super-resolution reconstruction

The single-molecule localization and reconstruction were performed on ThunderStorm^27^ based on ImageJ^28^. Wavelet filter (B-Spline) was exploited for the identification of potential molecular signals. Area of 7 × 7 pixels was extracted for each single-molecule candidate and further fitted with an integrated form of 2D gaussian function through a maximum likelihood approach. Lateral shifts during the imaging were corrected through a cross-correlation analysis. Localization with either large/small widths (> 1.5 × median(sigma) & < 0.5 × median(sigma)) or low brightness (< 100 photons) were considered as noise signals and removed for further analysis. Super-resolution images were reconstructed by an “averaged shifted histogram” method at a 10-fold magnification.

### Single-molecule tracking

The localization (from the reconstruction) were directly utilized for single-molecule tracking analyses. To avoid the merging or splitting events from high-density single-molecule signals during the initial acquisition, localization from 6000-9000 frames were selected. According to the diffusion model (equation (3)),^29^ molecules with diffusion coefficients of 0.1-0.5 μm^2^/s have probabilities of 0.91-1 for reappearing within 0.312 μm (the width of 3 pixels) distance on the consequent acquisition frame at a interval time of 20 ms. Therefore, 3 pixel was set as the upper searching bound for linking during our tracking analyses on U-tracker^30^.

### Post-analysis on dynamics

Multi-level analyses were conducted to study the diffusion of BDP-Mem (> 10 steps) by a home-written Matlab software.

I. Analysis of single trajectory. The diffusive pattern of each molecule was studied. The square displacement (*r*^2^) of diffusion at time t after time lag Δt (frame acquisition time, 20 ms in this study) is defined according equation (1).

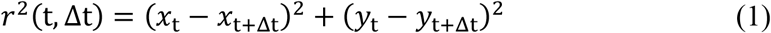

The mean square displacement (MSD) of diffusion after time lag Δt was calculated according to equation (2). N is the number of time lags (Δt) within a track.

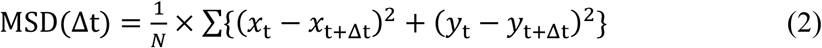

The cumulative density distribution (CDF) of square displacements was analyzed and fitted with a single-exponential function (equation (3)) in most cases, to calculate the diffusion coefficient (D) of a molecule.

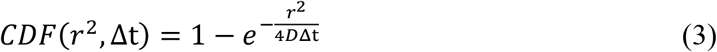

For a few molecules, they exhibited large variances on velocities during diffusion, and the CDF of their square displacements were fitted with a double-exponential function (equation (4)). In this case, the diffusion coefficient of a molecule was calculated following equation (5).

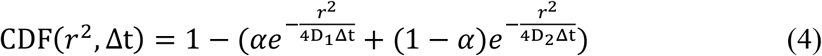

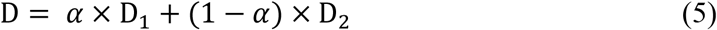

with α as the proportion of D_1_.

II. Analysis of single cell dynamics. Dynamics of living RBC membranes at the ensemble level was an average of the molecular diffusion distribution at the singlemolecule level. Thus, membrane dynamics could be quantitatively evaluated by an average diffusion coefficient 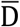 of BDP-Mem on the RBC membranes. 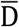 was obtained through statistical analyzing the probability density function (PDF) of the diffusion coefficient distribution on the same cell. For most cases, the PDF was fitted with a log-normal distribution,

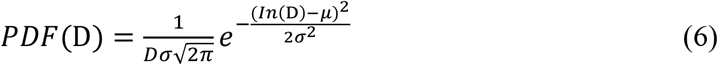

with μ as the mean and σ as the standard deviation of the log-normal distribution. The average diffusion coefficient 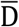 was obtained as,

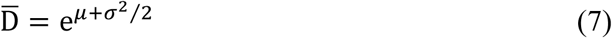

If motions of molecules on the cell showed both free and confined diffusions, a double log-normal distribution model was utilized for fitting the PDF,

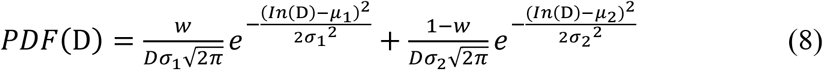

with μ_1_, μ_2_ as the means and σ_1_, σ_2_ as the standard deviation of two log-normal distributions. *w* is the proportion for the first distribution. For this case, the average diffusion coefficient (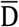) was calculated as:

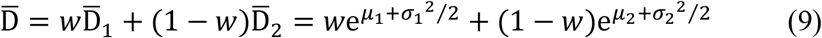

The transient velocities of probes on the membrane were also analyzed. The transient velocity is defined as the diffusion speed of a molecule between consequent steps on its trajectory.

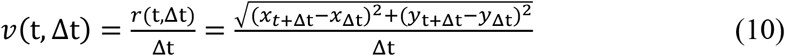

The diffusive velocity and direction of probes were mapped on the membranes. Since molecular motion of our probes reflects the membrane dynamics within nearby regions, spatially approximate measurements were utilized for determining the membrane features through convoluting with a weight function (a normalized gaussian function of 9 × 9 pixels size and 5 pixel sigma), which could avoid biases induced from single samplings.

### Synthesis of a far-red membrane probe, BDP-Mem

The π conjugation system of BODIPY was extended with two *p*-piperazinyl styryl groups through Knoevenagel condensation under inert atmosphere. The reaction byproduct water was continually removed from the mixture to proceed the condensation. The acquired product was purified and further quaternizated on its non-aryl amino moieties to obtain BDP-Mem. The details and characterizations of these syntheses are provided in section 6 and 7 of Supplementary Information.

### Fabrication of the microfluidic device

The mold of microfluidic devices was fabricated using two-layer lithography, which contained RBC microchamber structures with a height of 2 μm and main channel structures with a height of 80 μm. Polydimetylsiloxane (PDMS) and cross-linker (Sylgard 184, Dow Corning) were mixed at 9:1 ratio and poured on the mold, first degassed and cured at 80 °C for 24 h. Cured PDMS was peeled off from the mold and holes of the inlets were punched. The PDMS replica was covalently bound to a cell culture dish (CellVis, D29-20-1-N) after surface activation for 15 s in oxygen plasma (Diener electronic, GmbH, Germany). To avoid possible damages of cell membrane induced by the strong hydrophilic stress, the microfluidic device was utilized 1 hour after the oxygen plasma bounding.

### RBCs Staining

The study was performed with the approval from an institutional ethics committee of Dalian University of technology (#2021-054). Informed consents were provided by the blood donors. Fingertip blood was instantly mixed with an Alsever’s Solution (R1016, Solarbio) in a volume ratio of 1:1 to avoid coagulation. A stock solution of membrane probes was properly diluted with an isotonic buffer before staining. Concentrations utilized for staining included: (1) 5 μM BDP-Mem or 5 μM DiD for confocal comparison imaging (Fig.1c, Supplementary Figs.5-6); (2) 50 nM ~ 300 nM BDP-Mem for super-resolution imaging studies (Figs.2–4); (3) 200 nM BDP-Mem or 200 nM DiD for super-resolution comparison studies (Figs.2f-g). All staining was proceeded at room temperature for 5 min. Stained RBCs were washed with isotonic buffers for 3 times. A RBC suspension solution was obtained by concentration at 800 rpm/min for 2 min, and was utilized for the microfluidic device loading.

### Physiological condition for imaging

The microenvironments for RBCs were continual regulated by syringe pumps (Lange, China) at a constant rate 1 μL/min. An isotonic buffer (RBC buffer*+ 0.1% FBS) was utilized for preserving the normal living state of RBCs. Two classes of buffers were utilized to construct pathological conditions. (1) Osmotic stress buffers: (a) RPMI1640, supplemented with 10% FBS, water (50% vol%, 143 mOsm/kg, for ghosts; 15% vol%, 251 mOsm/kg, for stomatocytes); (b) RPMI1640, supplemented with 10% FBS, NaCl (0.5% wt%, 381 mOsm/kg; 1.0% wt%, 466 mOsm/kg) for echinocytes. (2) Pb^2+^ poisoning condition buffers: (c) RBC buffer, supplemented with 0.1% FBS, 10 μM Pb^2+^; (d) RBC buffer, supplemented with 0.1% FBS, 10 μM Pb^2+^, 1 μM Senicapoc; (e) RBC buffer, supplemented with 0.1% FBS, 1 μM Senicapoc. Under continual buffering conditions, deformed RBCs were imaged at corresponding time in the main text.

*The recipe of isotonic RBC buffer: 32 mM HEPES, 125 mM NaCl, 1 mM MgSO_4_, 1 mM CaCl_2_, 5 mM KCl and 5 mM Glucose, with pH adjusted to 7.2 (310 mOsm/kg).

### Single-molecule photophysical study

Cover slips (Fisherbrand, 12-545-102) were cleaned by sequential sonication in ethanol, 1 M KOH, MilliQ water, and dried by high-purity N_2_. Remaining fluorescent dusts or particles were removed through oxygen plasma treatment (Diener electronic, GmbH, Germany) for 3 min. BDP-Mem was diluted to 10 pM in 0.1% polyvinyl alcohol (PVA) aqueous solution to avoid molecular overlaps. Thin polymer films were obtained by spin-coating 200 μL diluted BDP-Mem PVA solution on the cover slip surface at 3000 rpm/min for 30 s. Dry films were obtained under low relative humidity (<10% RH) and wet films were acquired under high relative humidity (est. > 80% RH). Single-molecule analyses were conducted on the same TIRF microscopy as described in the section of super-resolution imaging (under the convention imaging mode). The frame acquisition time was 40 ms and 3000 frames were collected under each conditions. The raw stack images were analyzed by a home-written Matlab software as described previously.^16,26^ During the analysis, the software automatically searched for single-molecule candidates, extracted fluorescent trajectories and identified the state-transforming events. On the basis of these data, the photophysics of single molecules were calculated following the methods in section 3 of Supplementary Information.

## Acknowledgements

This work is supported by National Natural Science Foundation of China (Nos. 21576040, 21776037, 21804016, 22004011 and 11974002), China Postdoctoral Science Foundation (No. BX20200073 and 2020M670754), the Fundamental Research Funds for the Central Universities (No. DUT20JC39) and Dalian Science and Technology Innovation Fund (No. 2020JJ25CY014). The fluorescent imaging is performed with support from Chemical Analysis and Research Center, Dalian University of Technology.

## Author contributions

Y. X. conceived the research. Z. Y. and W. Y. designed and conducted the experiments, analyzed the collected data, prepared the figures and wrote the paper. W. Y. and S. W. prepared the microfluidic chip. Y. Z. assisted in the analysis tool establishment. X. Z. and H. Y. synthesized the probe. C. L. designed the microfluidic chip. Y. X., W. Y., X. P. and C. L. revised the paper. All authors read and approved the final manuscript.

## Competing interests

The authors declare no competing interests.

## Notes

### Competing Interest Statement

The authors have declared no competing interest.

