## Supplementary Information for "Imaging the ultrastructures and dynamics of live erythrocyte membranes at the single-molecule level with a far-red probe on a microfluidic platform"

### 1 The challenges for imaging living erythrocytes

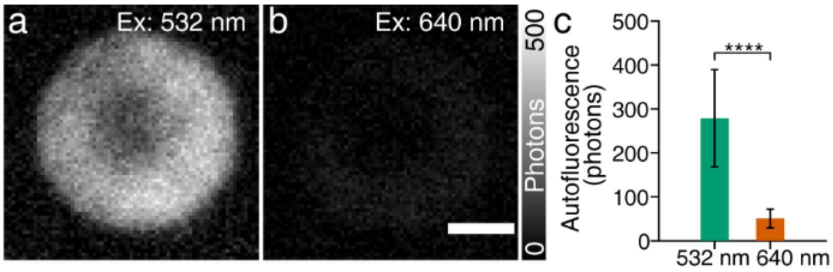

Supplementary Fig.1. Autofluorescence of RBCs. The fluorescence from a RBC excited under a 532 nm (a) or 640 nm (b) laser. The powers of two excitation lasers were adjusted to be identical. (c) Comparison on the average autofluorescence of RBCs when excited at different wavelengths. Significance was determined through a two-sample t-test with $P < 0.0001$ **** (sample size n = 214 RBCs). Hemoglobin-rich RBCs demonstrate intensive visible absorption.[1,2] When excited with a visible-light laser (e.g. 532 nm), this feature results in potential photodamage and significant background autofluorescence (a), the latter of which further severely blurs the super-resolution reconstruction. The utilization of a red-shifted laser at 640 nm would significantly reduce the autofluorescence, resulting in enhanced signal-to-noise ratio beneficial for imaging. Error bars indicate the s.d.. Scale bar: 2 µm.

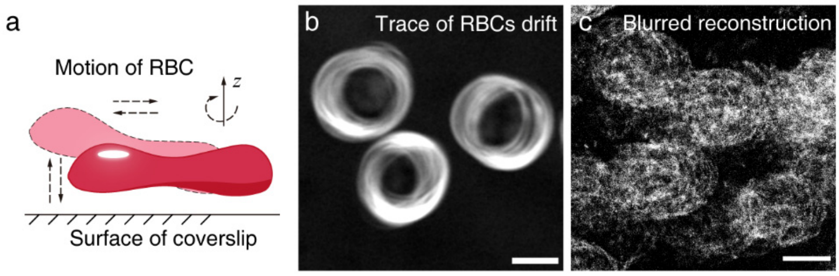

Supplementary Fig.2. Motion of RBCs challenges the super-resolution imaging of their membranes. (a) Illustrative chart presents the probable motion of RBCs during imaging acquisition. (b) Overlaid edges of RBCs within 1.5 min in a static buffer. (c) Blurred super-resolution reconstruction of RBC membranes due to motion. RBCs are restless during imaging, due to the flow of culture media or the unbalanced molecular collisions. Even if settled in static buffer, living RBCs are easily movable during imaging, contradicted to the enduring accumulation required by super-resolution imaging. Above observation is in agreement with previous results, which utilizes chemical fixation method[3,4]. The irregular drift ultimately leads to the blur of super-resolution reconstruction. Scale bars: 5 µm.

#### 2 Spectral studies

Spectral measurement: Fluorophores were dissolved in dimethylsulfoxide to prepare 1 mM stock solution. Before measurement, sample solution was obtained through diluting the stock solution to a desired concentration in target solvents (typical 1 μM). The volume of dimethylsulfoxide in the target solution was ≤ 0.1%. Samples were transferred to quartz cuvettes (4 mL) for measuring. Absorption spectra (precision: ± 1.5 nm) were recorded on Agilent 8453 UV-Vis spectrophotometer. Fluorescence spectra (precision: ± 1.5 nm) were recorded on Agilent Cary Eclipse Fluorescence spectrophotometer.

The absolute quantum yield of BDP-Mem in dichloromethane was determined through an absolute photoluminescence quantum yield spectrometer (Hamamatsu C11347). This solution of BDP-Mem in dichloromethane was further employed as the reference standard to determine the relative quantum yield (precision: ± 10%) of BDP-Mem in other solvents. The relative quantum yield was determined according to the equation (1):

$$\Phi_s = \Phi_{st} \times \frac{A_{st}}{A_s} \times \frac{I_s}{I_{st}} \times \frac{n_s^2}{n_{st}^2} \tag{1}$$

$\Phi_s$ was the relative quantum yield of a sample. $\Phi_{st}$ was the quantum yield of the standard. $A_s$ and $A_{st}$ were the absorption of the sample and standard at the excitation wavelength. $I_s$ and $I_{st}$ were the fluorescence integration of sample and standard, respectively. $n_s$ and $n_{st}$ were the refractive index of the utilized solvents.

RBC ghosts: Blood was collected in venous blood tubes contained EDTA-$K_2$. Fresh blood was centrifugated at 2000 g for 30 min at room temperature and the supernatant (plasma and buffy coat layer) was discarded. Then the acquired RBCs were mixed with Alsevier solution in a volume ratio of 1:1 for storage at 4 °C. Before extraction, these stored RBCs were mixed with PBS in a ratio of 1:5 and centrifugated at 2000 g for 5 min to remove lysed cells. Normal RBCs were further diluted in PBS to achieve a volume of 1 mL. To hydrolysis RBCs, the osmolarity of this RBC suspension buffer was adjusted to ~ 20 imOsm[36] through the addition of 14 mL pre-diluted PBS buffer. This hydrolysis process was proceeded for 40 min at 4 °C. After total hydrolysis of RBCs, the ghosts were obtained through centrifugation at 20000 g. These ghosts were washed for three times with diluted PBS buffers at identical osmolarity (~20 imOsm) to remove excess hemoglobin. Finally, these ghosts were transferred to 200 μL PBS to recover their morphology. The obtained ghosts PBS solution was stored in refrigerator at 4 ºC.

Ghosts staining: Fluorophore stock solution was diluted to prepare 1 μM probe PBS buffer at a total volumn of 3 mL. 10 μL ghost PBS solution was added to this probe solution in 4 mL centrifugation tubes. The solution was instantly mixed through several times of upside-down. The staining was proceeded on a stage shaker (Beyotime) at a speed of 120-140 rpm/min for 1 h before measurement.

Supplementary Table 1. Spectral properties of BDP-Mem in various solvents

| Solvent | $\lambda_{ab}$ (nm) | $\lambda_{em}$ (nm) | $\varepsilon \times 10^4$ ($mol \cdot L^{-1} \cdot cm^{-1}$) | Φ |
|---|---|---|---|---|
| DCM | 650 | 671 | 10.0 | 0.52 |
| Methanol | 652 | 681 | 9.8 | 0.54 |
| Water | 651 | 711 | 6.9 | 0.05 |
| DMF | 669 | 711 | 8.4 | 0.32 |
| DMSO | 677 | 727 | 8.9 | 0.21 |

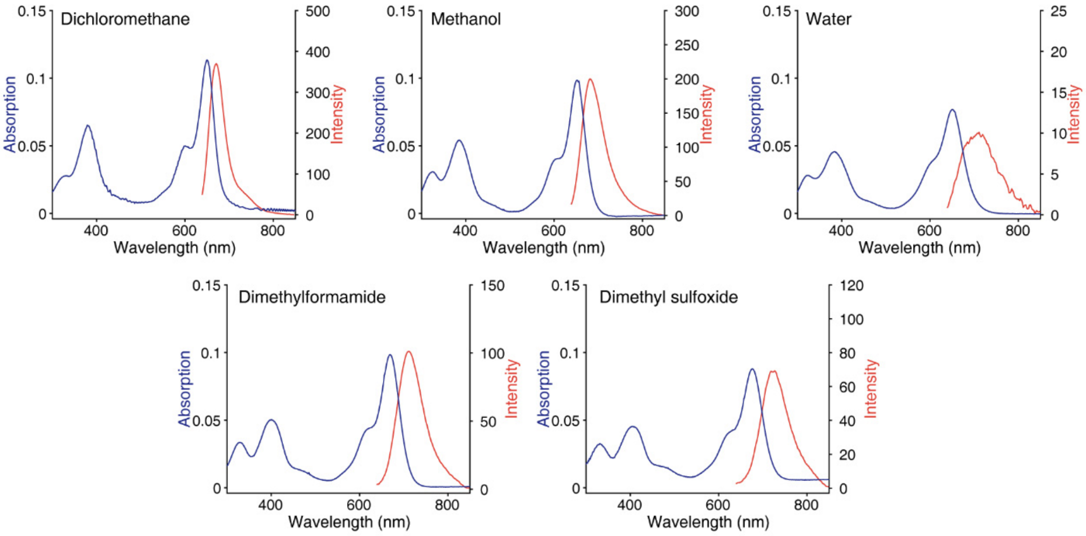

Supplementary Fig.3. Absorption and fluorescence spectra of BDP-Mem in various solvents.

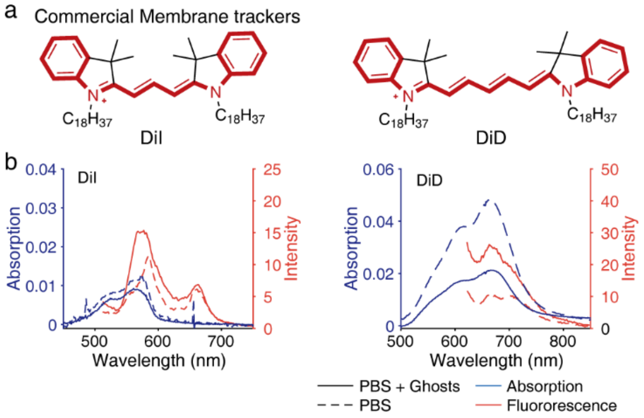

Supplementary Fig.4. Spectral changes of commercial membrane probes upon staining RBC ghosts. (a) Molecular structures of commercial membrane probes, DiI and DiD. (b) Absorption and fluorescence spectra of those probes upon staining RBC ghosts.

#### 3 Single-molecule analysis

**Photon conversion.** The intensity from each pixel ($I_{pixel}$) was converted to photons through $\text{Photons} = (I_{pixel} \times \text{ADU})/(\text{EMGain} \times \text{QE})$. ADU was the CCD sensitivity (15.82 electrons per A/D counts) obtained from the performance sheet of the EMCCD camera. EMGain was the gain configuration (100) for single-molecule imaging. QE (90-95) was the quantum yield of the camera sensor.

**Single-molecule brightness.** A single-molecule signal is composed by several pixels due to the light diffraction. The single-molecule brightness ($I_{sm}$) was defined as the photon collection from a single-molecule signal during one frame acquisition time (40 ms). It was calculated according to equation 2.

$$I_{sm} = \sum_{k \in K} (I_{pixel}^{k} - I_{background}) \tag{2}$$

$I_{pixel}^{k}$ was the photon number of pixel k; K was a set of pixels, which forms one single-molecule signal; $I_{background}$ was the background noise. The reported single-molecule brightness of BDP-Mem under dry or wet condition ($\bar{I}_{sm}^{condition}$) was a statistical average,

$$\bar{I}_{sm}^{condition} = \frac{\sum_{n=1}^{N_{signals}} I_{sm}^{n}}{N_{signals}} \tag{3}$$

with $N_{signals}$ as the total number of bright signals ($N_{signals} = \sum_{f \in F} N'_{f}$, with $N'_{f}$ as the number of signals on frame f, and F as a set of acquisition frames).

**Photons per switching cycle.** This parameter was calculated as the photons collection during a bright state:

$$Photons\ per\ switching\ cycle = \sum_{j \in J} I_{sm}^{j} \tag{4}$$

with $I_{sm}^{j}$ as the single-molecule brightness at frame j, and J as the set of consecutive frames from a bright state. The reported photons per switching cycle was a statistical average from all bright states.

**On time ($t_{on}$).** The time duration of a bright state. Reported on time was calculated as a statistical average,

$$\bar{t}_{on} = \frac{\sum_{n=1}^{N_{bs}} t_{on}^{n}}{N_{bs}} \tag{5}$$

with $t_{on}^{n}$ as the on time of bright state *n*, and $N_{bs}$ as the total number of bright states.

**Total collected photons before photobleaching.** This parameter was calculated as the sum of photon collections from bright states of a molecule,

$$Total\ collected\ photons = \sum_{l \in L} \mathrm{I}_{sm}^{l} \tag{6}$$

with $\mathrm{I}_{sm}^{l}$ as the single-molecule brightness at frame *l*, and L as a set of frames from bright states of a molecule. The reported value was a statistical average from molecules under the same condition.

**The number of Photoswitching cycles.** The number of bright-to-dark switches happened for a fluorophore molecule before photobleaching. The reported value ($\overline{N}_{cycles}$) was a statistical average,

$$\overline{N}_{cycles} = \frac{\sum_{n=1}^{N_m} N_{cycles}^{n}}{N_m} \tag{7}$$

with $N_{cycles}^{n}$ as the number of photoswitching cycles from molecule *n*, and $N_m$ was the total number of investigated molecules.

### 4 Imaging & comparison studies

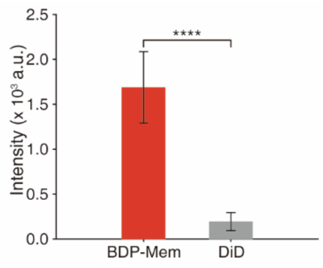

Supplementary Fig.5. Comparison on the confocal fluorescence imaging of BDP-Mem and DiD on living RBC membranes. Confocal images were collected on an inverted FV1000 microscope (Olympus, Japan) with a 60× (PLAPON) objective. The wavelength selection for both fluorophores was 655-755 nm. The difference significance was determined through a two-sample t-test $P<0.0001$, **** ($n > 50$ RBCs).

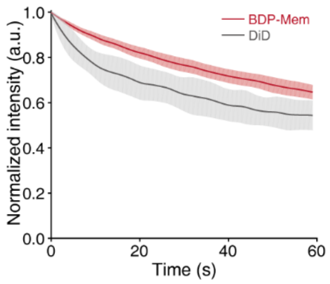

Supplementary Fig.6. Comparison on the photostability between BDP-Mem and DiD on the RBC membranes. RBCs were separately stained with 5 μM of each probe for 5 min and then transferred to our microfluidic chip. Laser light (640 nm) was continually irradiated at the RBCs at the TIRF mode. The fluorescent signals from the RBCs were recorded consistently on an EMCCD camera. (sample sizes: > 10 RBCs)

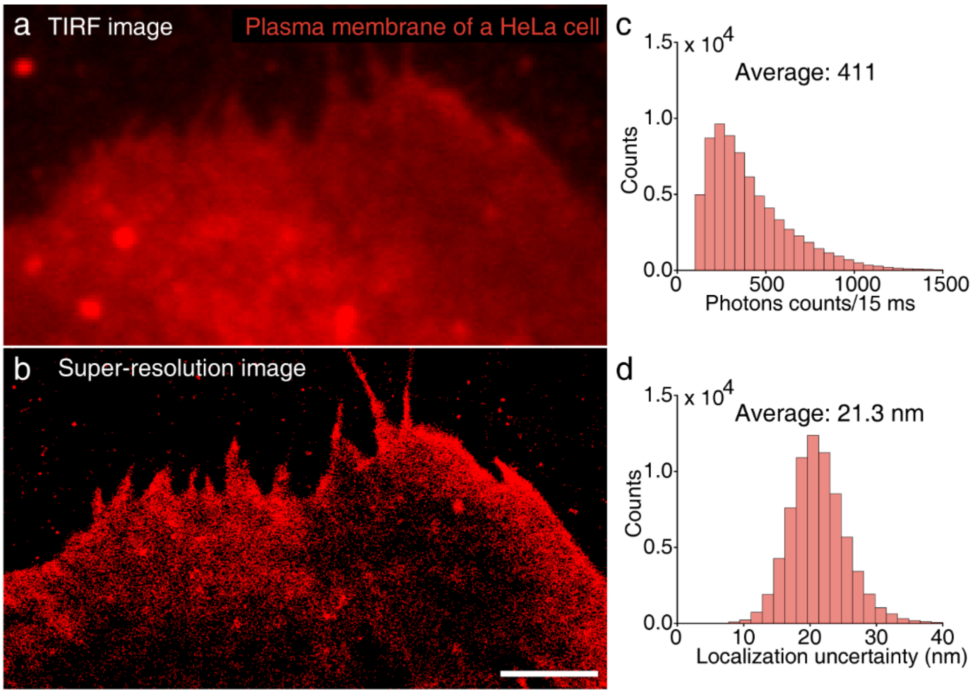

Supplementary Fig.7. Super-resolution imaging of living HeLa cell plasma membrane by BDP-Mem staining. (a) Conventional TIRF and (b) super-resolution reconstructed images of the plasma membrane. Histogram of single-molecule brightness (c) and localization uncertainties (d) of the reconstruction. Scale bar: 3 μm.

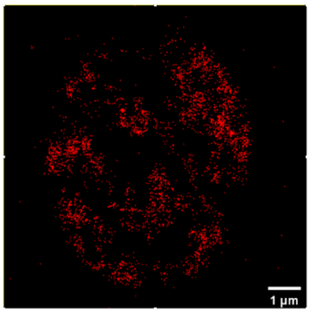

Supplementary Fig.8. Super-resolution reconstruction of RBC membrane through commercial probe DiD under the TIRF mode.

### 5 High-throughput imaging results

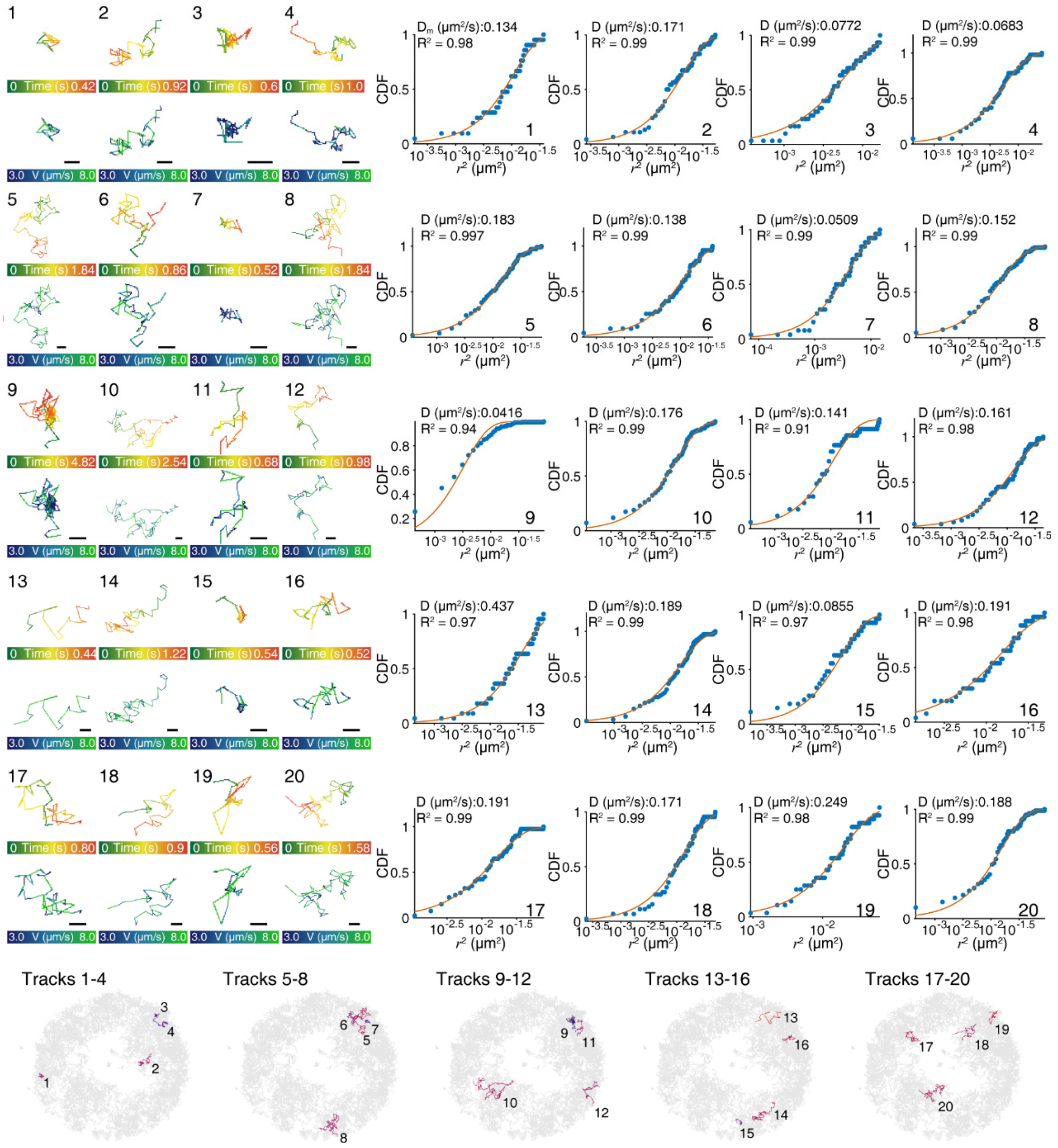

Supplementary Fig.9. Trajectories of BDP-Mem molecules on a RBC membrane. For each track, the left column shows time and velocity trajectories; the right column presents the cumulative distribution function (CDF) of square displacements ($r^2$). The CDF was fitted with a simple exponential decay. The bottom row displays the track spatial positions on the membrane. Scale bars: 200 nm.

#### Cell 1-5: Normal red blood cells (310 mOsm/kg)

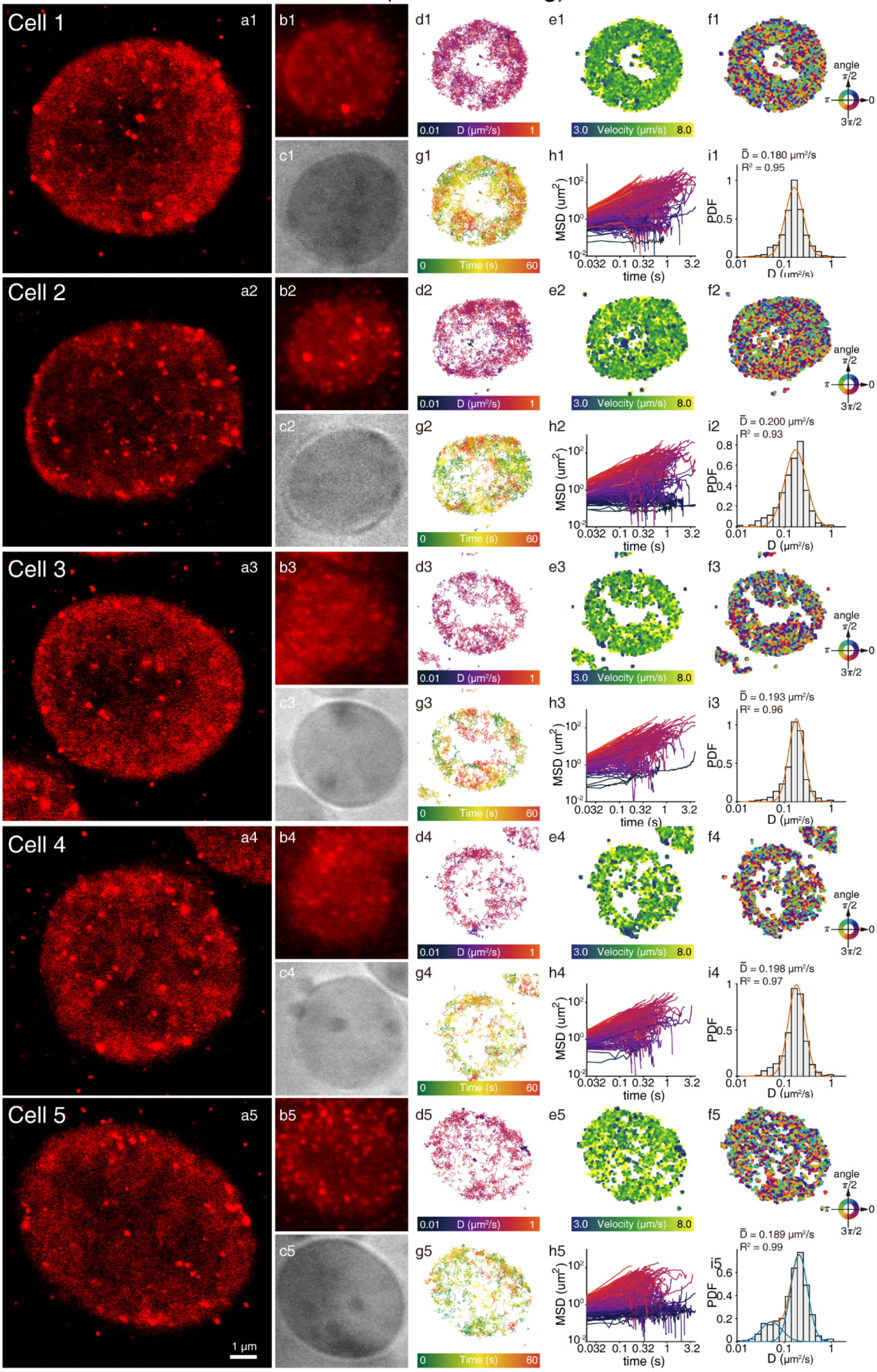

### Cell 6-10: Red blood cell "ghosts" (143 mOsm/kg)

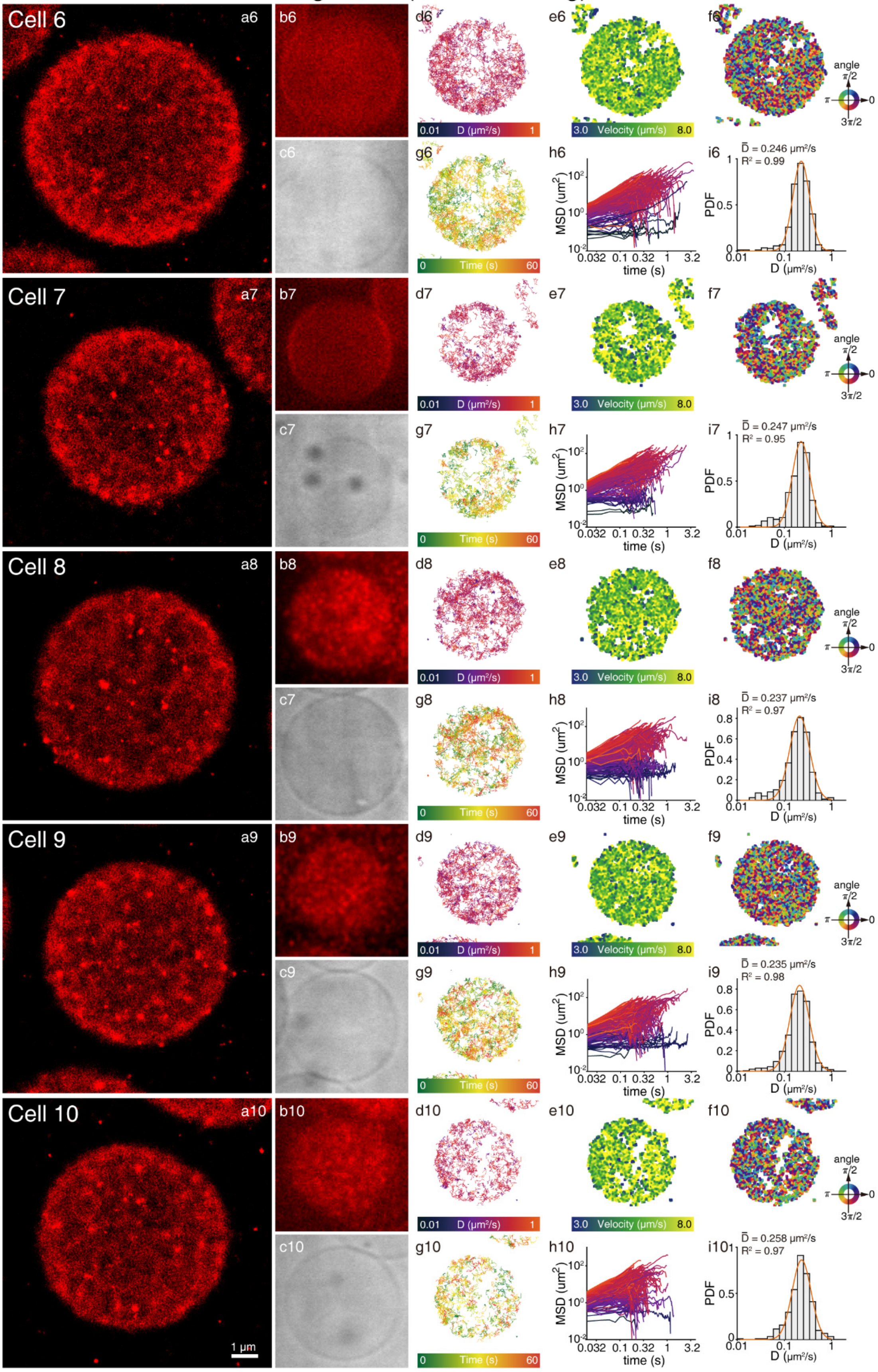

#### Cell 11-15: Stomatocytes (251 mOsm/kg)

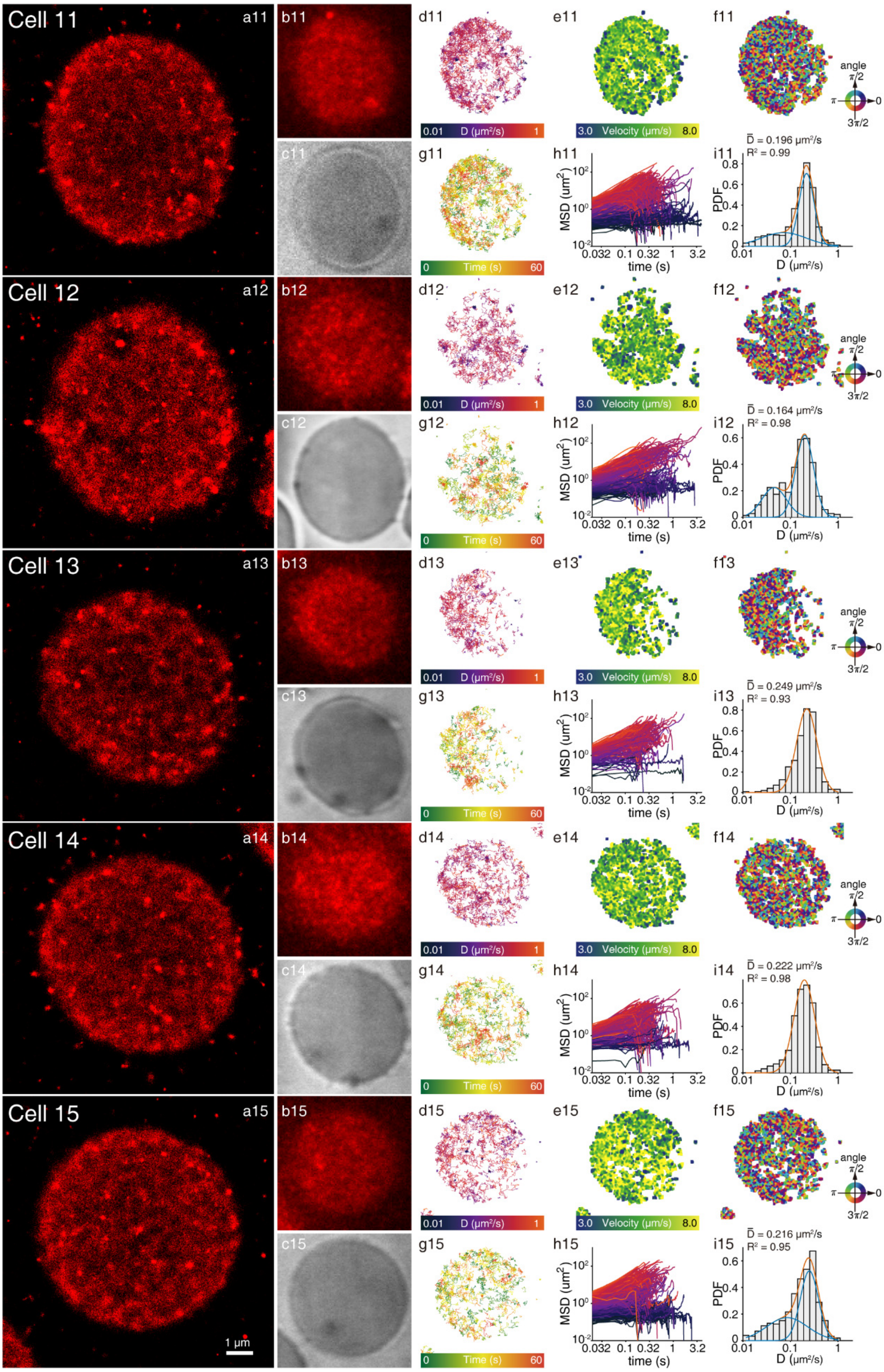

#### Cell 16-20: Echinocytes (381 mOsm/kg)

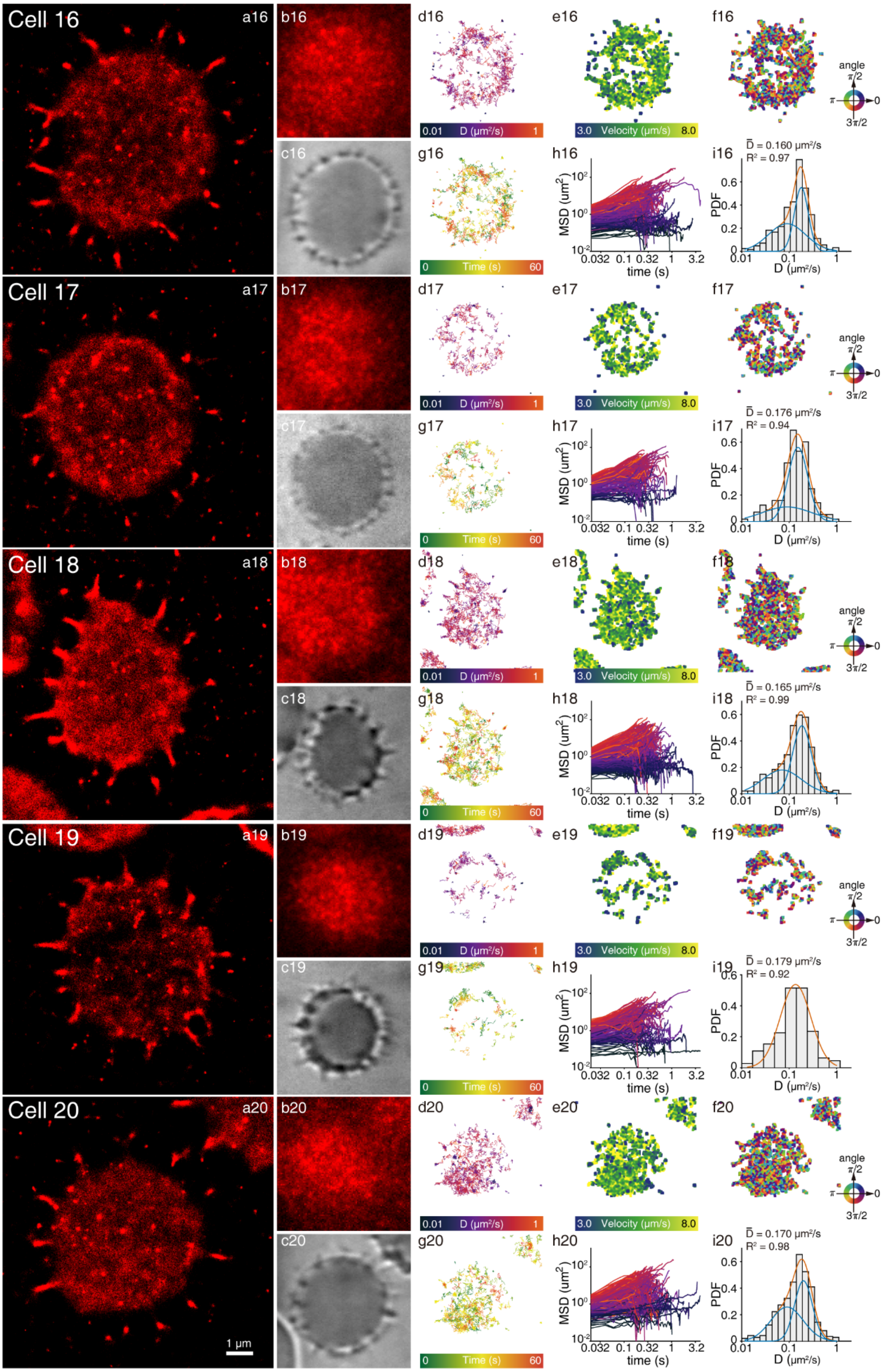

Cell 21-25: Echinocytes (466 mOsm/kg)

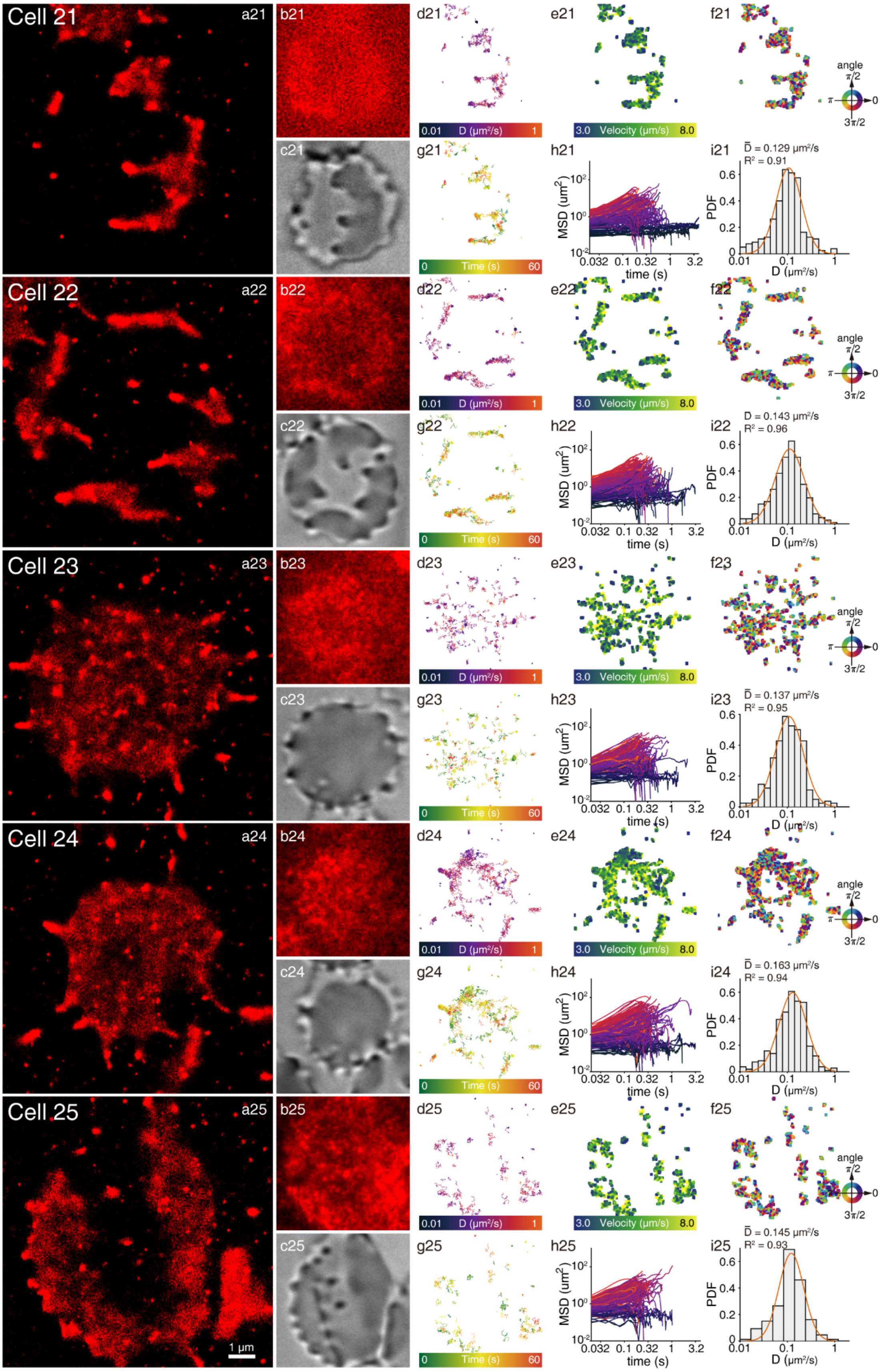

Cell 26-30: RBCs incubated with Senicapoc

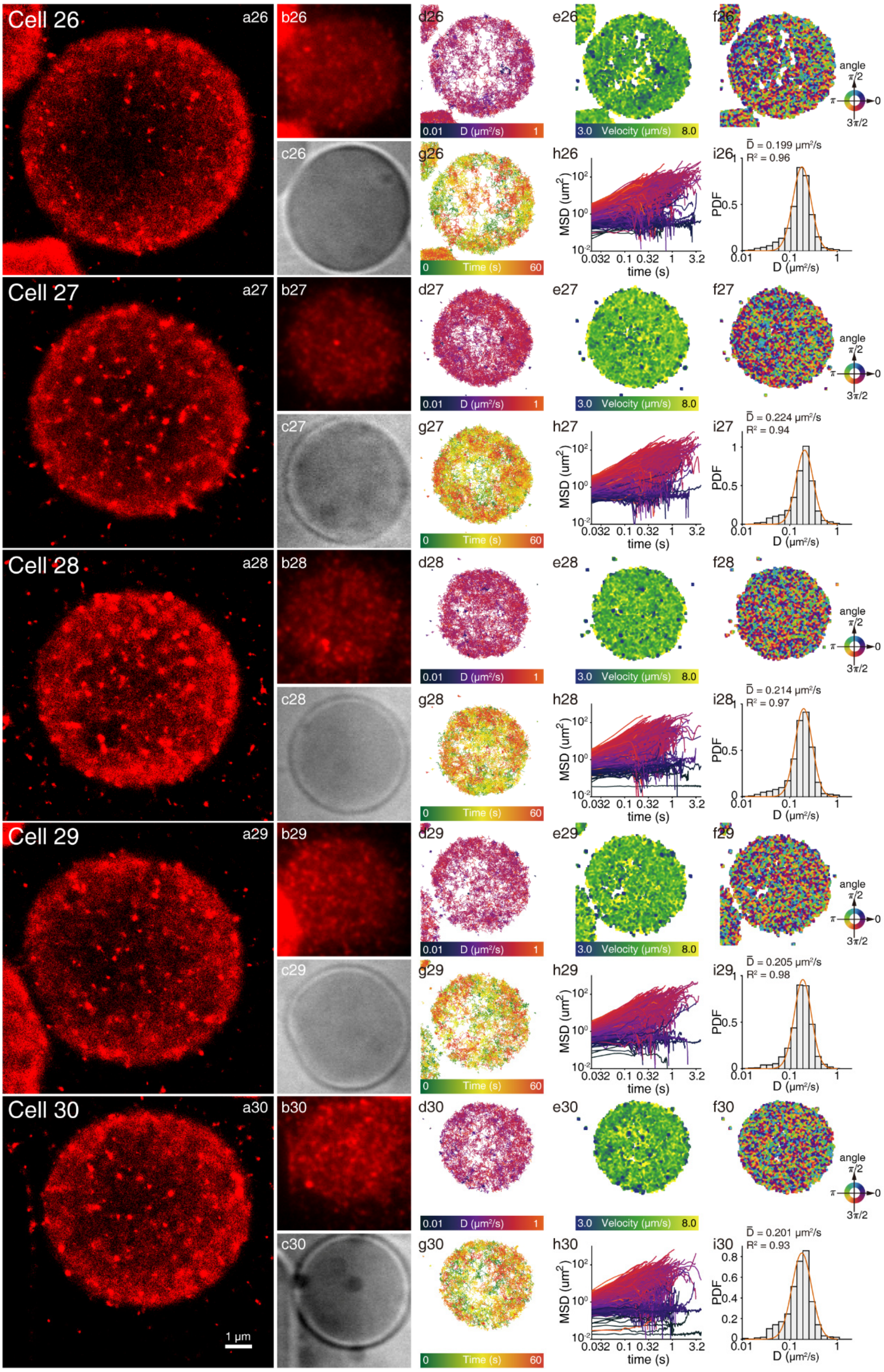

#### Cell 31-35: RBCs incubated with Senicapoc and $Pb^{2+}$ (t = 0-1 h)

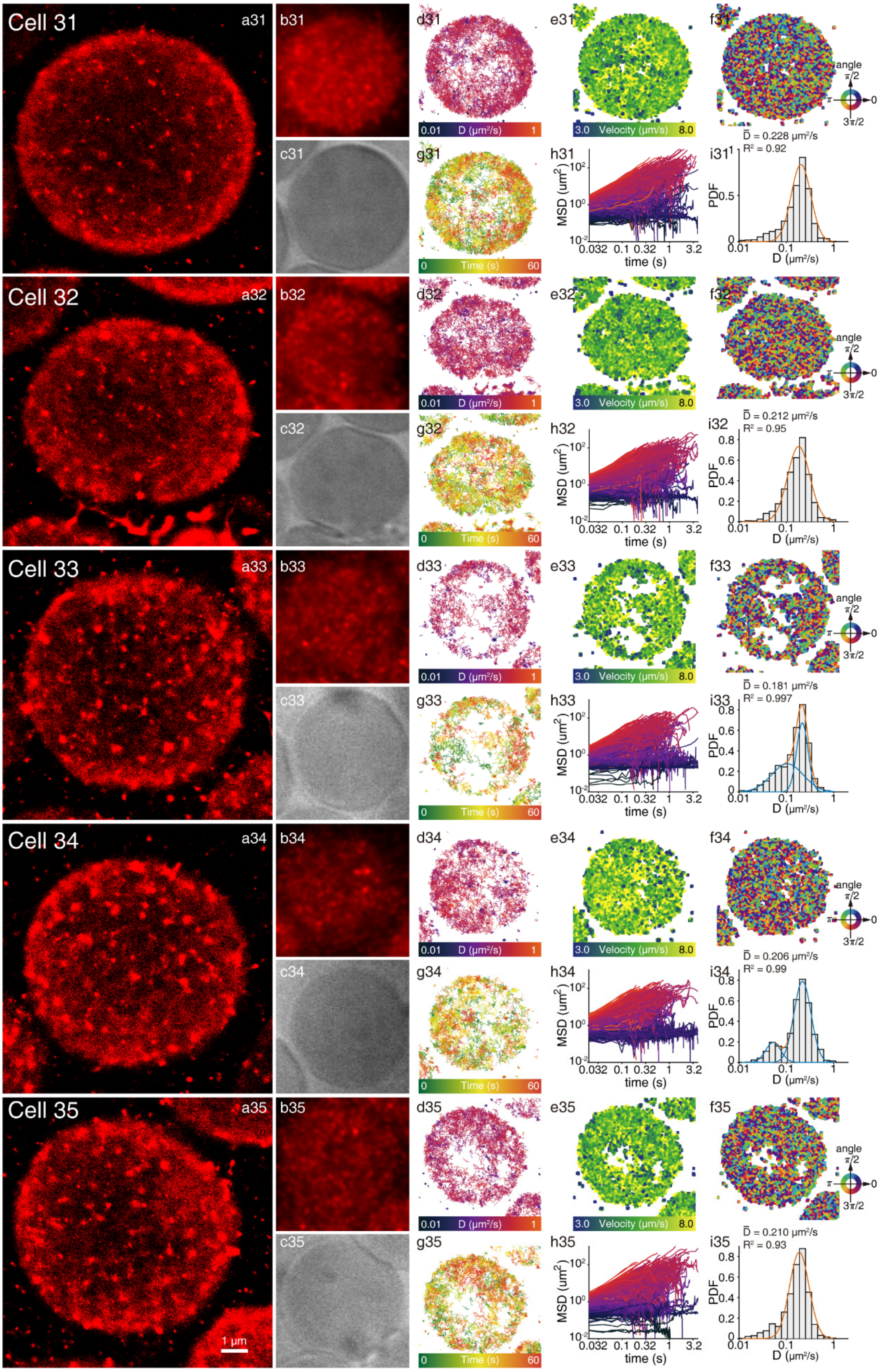

Cell 36-40: RBCs incubated with Senicapoc and $Pb^{2+}$ (t = 1-2 h)

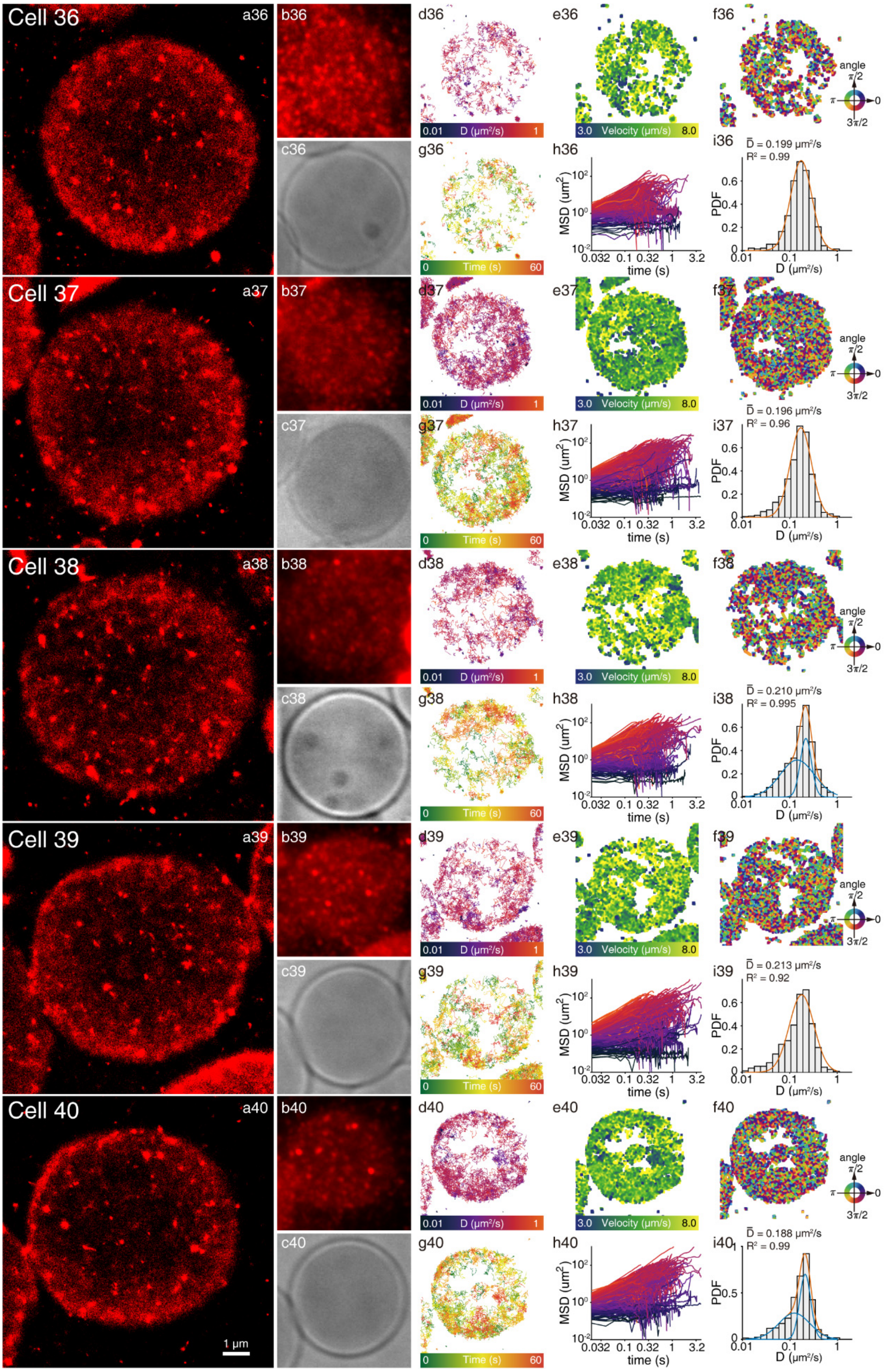

Cell 41-45: RBCs poisoned by $Pb^{2+}$ (t = 0-1 h)

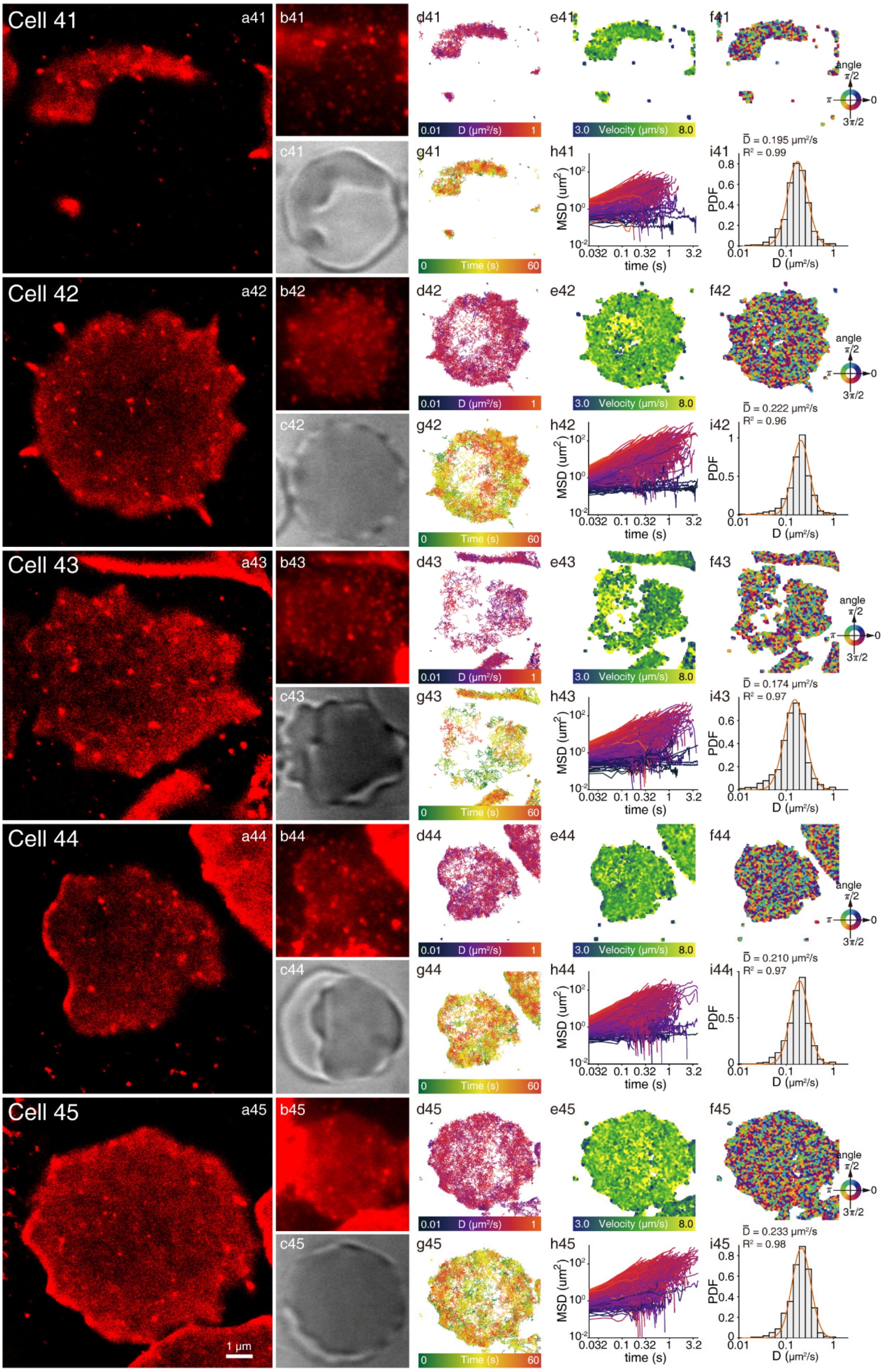

Cell 46-50: RBCs poisoned by $Pb^{2+}$ (t = 1-2 h)

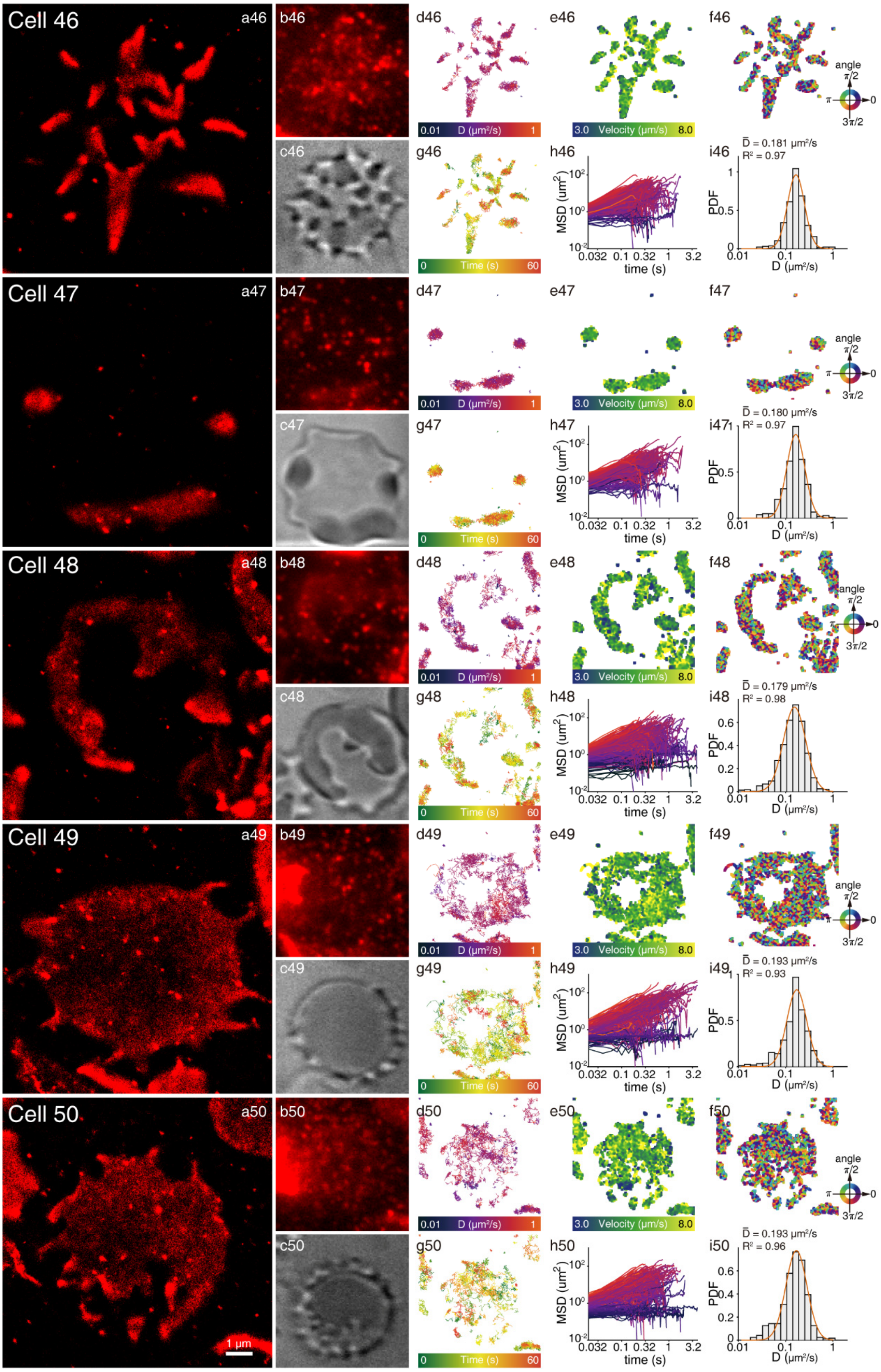

Supplementary Fig.10. High-throughput studies of the ultrastructures and dynamics of living RBC membranes. Living RBCs were stained with BDP-Mem and immobilized in our microfluidic chip. Through the manipulation of RBC physiological conditions in our on-chip study, 10 different RBC conditions were constructed for analyzing RBC ultra-details and quantitative information. For each cell: the left shows super-resolution reconstruction (a1-50), conventional TIRF fluorescent images (b1-50) and bright-field images (c1-50) of RBC membranes; the right shows: single-molecule trajectory overlays (d1-50), velocity maps (e1-50), direction maps (f1-50), time-colored trajectories (g1-50), MSD plots (h1-50) and the diffusion coefficient distributions (i1-50) of BDP-Mem on the same RBC membrane. Cells 1, 3, 10, 14, 16, 31, 39, 42 and 46 are the same cells from Figs.3-5.

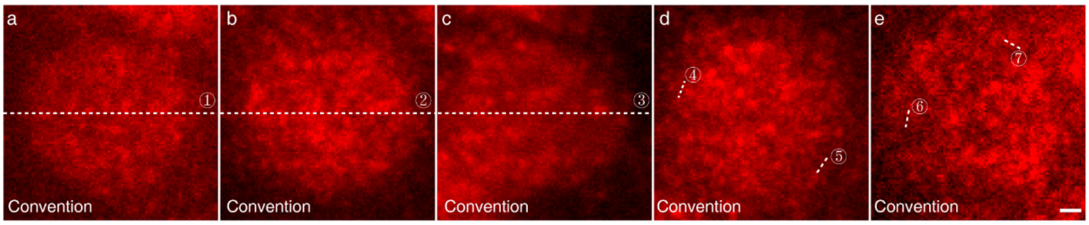

Supplementary Fig.11. Highlightened lines on conventional images of RBCs in Fig.4. Scale bar: 1 μm.

Supplementary Table 2. Difference significance analysis of the diffusion coefficients (Fig.5a) on the RBC membranes under osmotic stress.

| Cell Type A | Cell Type B | p | Significance |
|---|---|---|---|
| Ghosts | Stomatocytes | $1.49 \times 10^{-2}$ | * |
| Ghosts | Echinocytes (381 mOsm/kg) | $<1.00 \times 10^{-6}$ | **** |
| Ghosts | Echinocytes (466 mOsm/kg) | $<1.00 \times 10^{-6}$ | **** |
| Stomatocytes | Echinocytes (381 mOsm/kg) | $2.65 \times 10^{-2}$ | * |
| Stomatocytes | Echinocytes (466 mOsm/kg) | $3.00 \times 10^{-6}$ | **** |
| Echinocytes (381 mOsm/kg) | Echinocytes (466 mOsm/kg) | $6.22 \times 10^{-3}$ | ** |

Supplementary Table 3. Difference significance analysis of the diffusion coefficients (Fig.5b) on the RBC membranes under $Pb^{2+}$ poisoning conditions.

| Condition A | Condition B | p | Significance |
|---|---|---|---|
| Normal | $Pb^{2+}$ (0-1 h) | $9.43 \times 10^{-1}$ | Null |
| Normal | $Pb^{2+}$ (1-2 h) | $1.53 \times 10^{-1}$ | Null |
| Normal | Senicapoc + $Pb^{2+}$ (0-1 h) | $4.91 \times 10^{-1}$ | Null |
| Normal | Senicapoc + $Pb^{2+}$ (1-2 h) | $5.31 \times 10^{-1}$ | Null |
| Normal | Senicapoc | $1.09 \times 10^{-2}$ | * |
| $Pb^{2+}$ (0-1 h) | $Pb^{2+}$ (1-2 h) | $2.43 \times 10^{-1}$ | Null |
| Senicapoc + $Pb^{2+}$ (0-1 h) | Senicapoc + $Pb^{2+}$ (1-2 h) | $9.60 \times 10^{-1}$ | Null |
| $Pb^{2+}$ (0-1 h) | Senicapoc + $Pb^{2+}$ (0-1 h) | $6.47 \times 10^{-1}$ | Null |
| $Pb^{2+}$ (1-2 h) | Senicapoc + $Pb^{2+}$ (1-2 h) | $1.56 \times 10^{-2}$ | * |

#### 6 Synthetic method

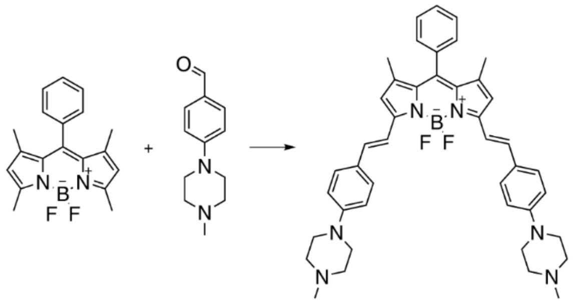

Synthesis of PBDP: BODIPY (500 mg, 1.54 mmol) and 4-(4-methylpiperazinyl)benzaldehyde (**1**, 788 mg, 3.86 mmol) were dissolved in a mixed solvents of toluene (50 mL) and piperidine (3 mL). An inert atmosphere was obtained through three times exchange of argon. After stirring the mixture for 30 min at room temperature, glacial acetic acid (3 mL) was added. The mixture was heated to reflux at a vigorous stirring condition. The byproduct water was continually removed from the system through a Dean-Stark trap. After refluxing for 56 h, the solution was concentrated in vacuo. The residue was purified by silica gel column chromatography using $CH_2Cl_2/CH_3OH$:15/1 as the eluent to obtain the 709 mg PBDP (yield: 66.1%). $^1$H NMR (500 MHz, $CDCl_3$) δ 7.61 (s, 1H), 7.58 (s, 1H), 7.54 (d, $J$ = 8.7 Hz, 4H), 7.51 – 7.44 (m, 3H), 7.32 (dd, $J$ = 7.1, 2.0 Hz, 2H), 7.20 (s, 1H), 7.17 (s, 1H), 6.91 (d, $J$ = 8.7 Hz, 4H), 6.60 (s, 2H), 3.31 (t, 8H), 2.58 (t, 8H), 2.36 (s, 6H), 1.42 (s, 6H). $^{13}$C NMR (101 MHz, $CDCl_3$) δ 152.79, 151.56, 141.45, 137.26, 136.05, 135.53, 133.20, 129.05, 128.99, 128.87, 128.75, 127.83, 117.53, 116.33, 115.31, 55.03, 48.27, 46.30, 14.73. HRMS (ESI) $m/z$ calcd for $C_{43}H_{49}BF_2N_6^{2+}$ $[M+2H]^{2+}$: 349.2034; found: 349.2036 ($z$ = 2).

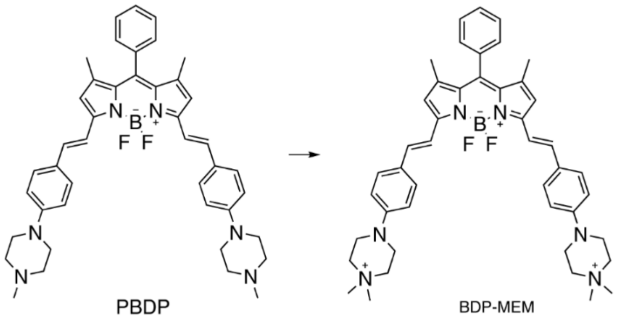

Synthesis of BDP-Mem: PBDP (50 mg, 71.8 μmol) was dissolved in 20 mL DMF, following the dropwise addition of methyl iodide (447 μL, 7.18 mmol). The reaction mixture was stirred at room temperature for 48 hours. After removing the solvent from the reaction system, the residue was washed with $CH_2Cl_2$ and n-hexane successively for three times respectively to offer 52 mg BDP-Mem (yield: 99.7%). $^1$H NMR (500 MHz, DMSO) δ 7.70 – 7.53 (m, 7H), 7.51 (d, $J$ = 16.1 Hz, 2H), 7.46 – 7.41 (m, 2H), 7.38 (d, $J$ = 16.2 Hz, 2H), 7.11 (d, $J$ = 8.3 Hz, 4H), 6.93 (s, 2H), 3.77 – 3.62 (m, 8H), 3.62 – 3.53 (m, 8H), 3.22 (s, 12H), 1.40 (s, 6H). HRMS (ESI) *m/z* calcd for $C_{45}H_{53}BF_2N_6^{2+}$ $[M]^{2+}$: 363.2191; found: 363.2187 ($z$ = 2).

### 7 Characterization spectra

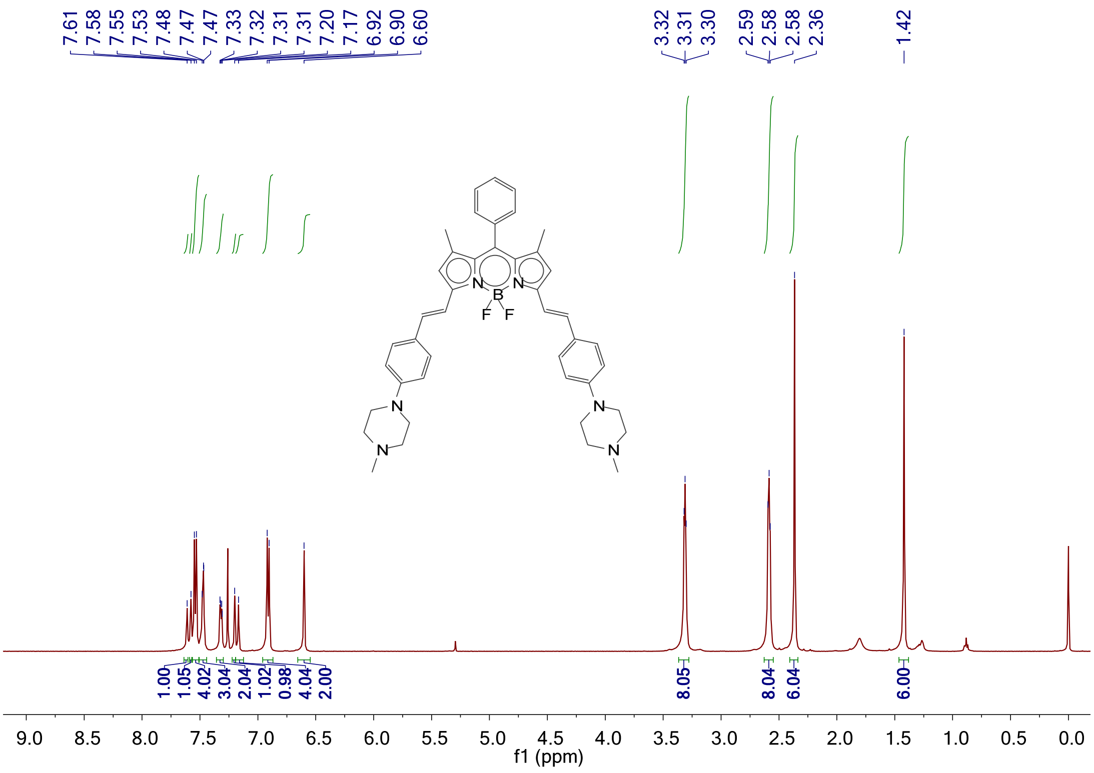

Supplementary Fig.12. $H^1$ NMR spectrum of PBDP.

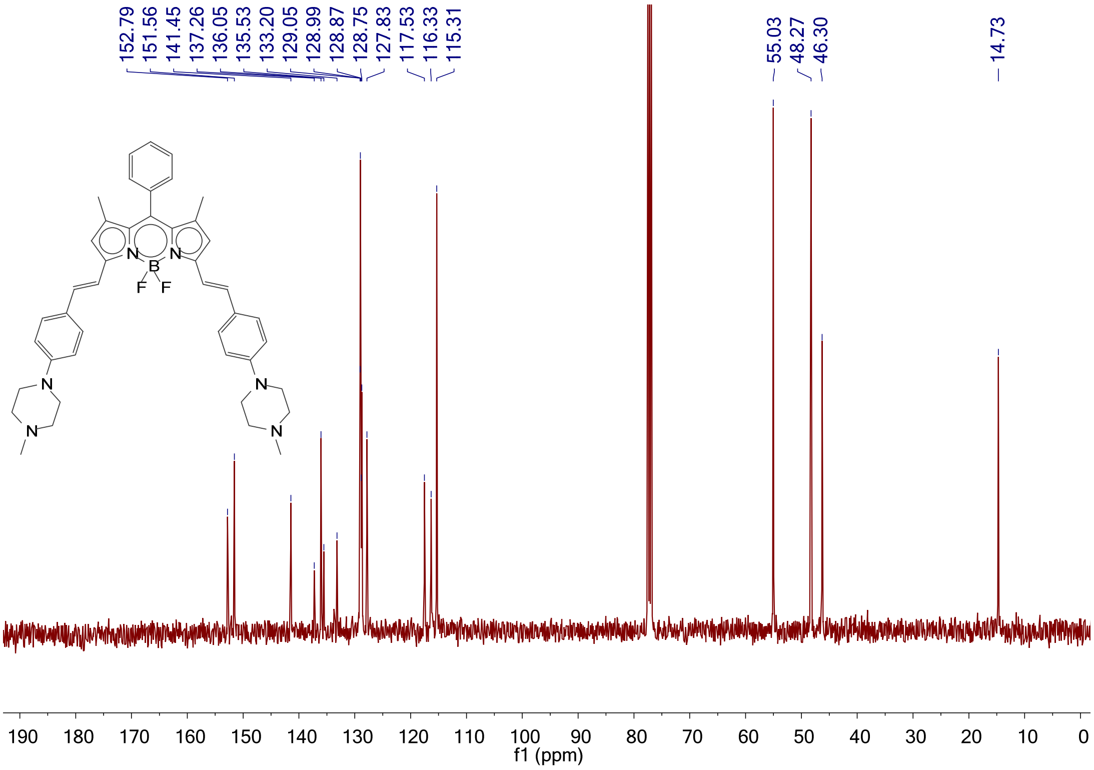

Supplementary Fig.13. $C^{13}$ NMR spectrum of PBDP.

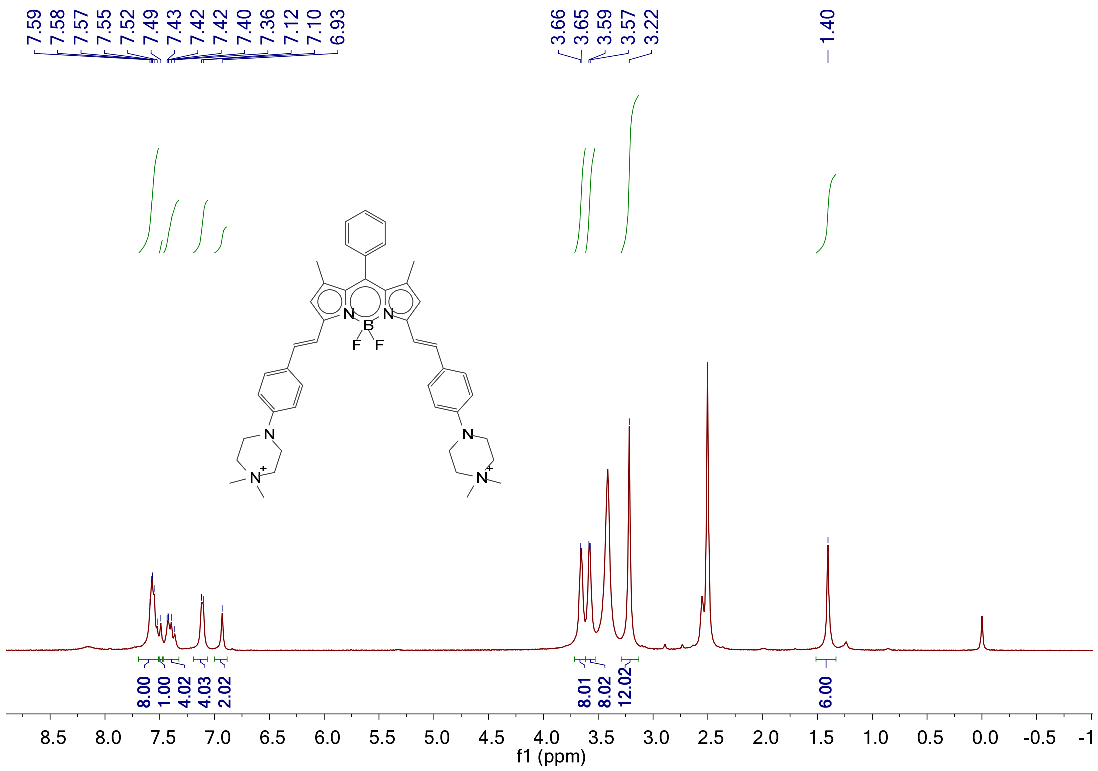

Supplementary Fig.14. H[1] NMR spectrum of BDP-Mem.

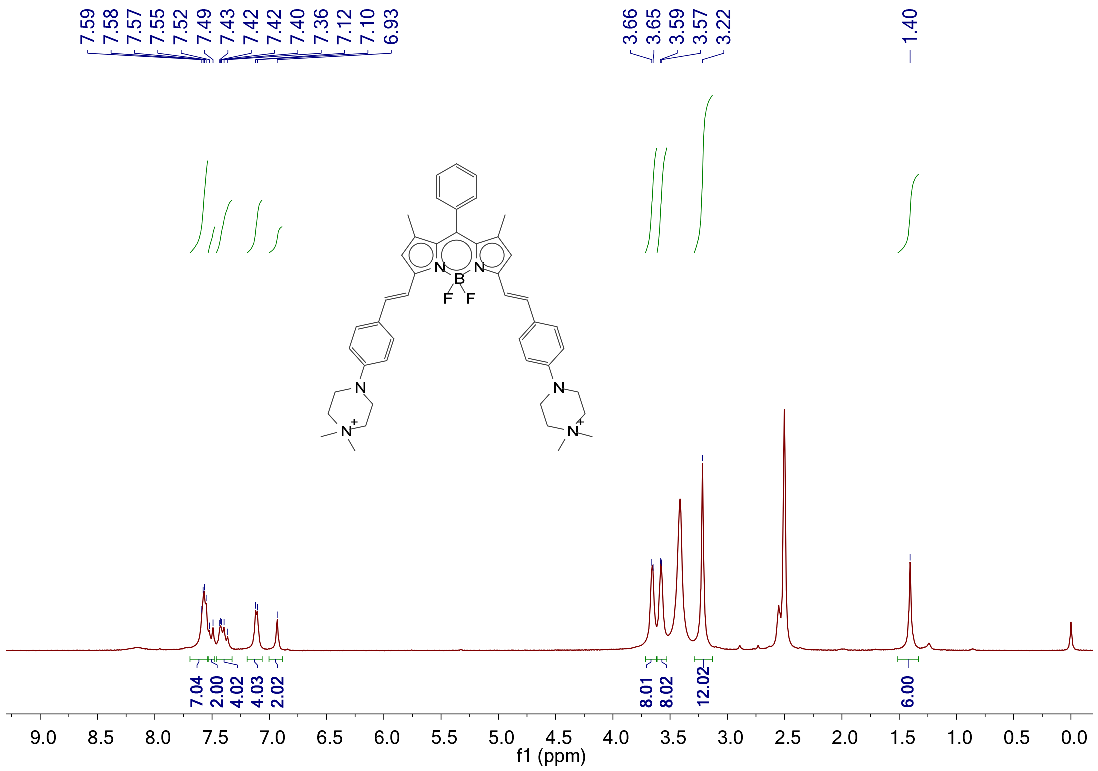

Supplementary Fig.15. C[13] NMR spectrum of BDP-Mem.
